# Defining the function of disease variants with CRISPR editing and multimodal single cell sequencing

**DOI:** 10.1101/2024.03.28.587175

**Authors:** Yuriy Baglaenko, Michelle Curtis, Majd Al Suqri, Ryan Agnew, Aparna Nathan, Hafsa M Mire, Annelise Yoo Mah-Som, David R. Liu, Gregory A. Newby, Soumya Raychaudhuri

## Abstract

Genetic studies have identified thousands of individual disease-associated non-coding alleles, but identification of the causal alleles and their functions remain critical bottlenecks. Even though CRISPR-Cas editing has enabled targeted modification of DNA, inefficient editing leads to heterogeneous outcomes across individual cells, limiting the ability to detect functional consequences of disease alleles. To overcome these challenges, we present a multi-omic single cell sequencing approach that directly identifies genomic DNA edits, assays the transcriptome, and measures cell surface protein expression. We apply this approach to investigate the effects of gene disruption, deletions in regulatory regions, and non-coding single nucleotide polymorphisms. We identify the specific effects of individual SNPs, including the state-specific effects of an *IL2RA* autoimmune variant in primary human T cells. Multimodal functional genomic single cell assays including DNA sequencing bridge a crucial gap in our understanding of complex human diseases by directly identifying causal variation in primary human cells.

## Main Text

Application of large scale genome-wide association studies has identified thousands of complex trait loci including in autoimmune, metabolic, and infectious diseases^1^. Computational strategies are sometimes effective at either fine-mapping statistical association signals to individual variants or prioritizing potentially functional variants^2,3^. These methods have been instrumental in investigating complex disease loci but fall short of accurately assigning function to non-coding variants which account for most complex disease alleles. Accurate functionalization requires genomic editing to introduce the allele in primary cells and assess downstream effects.

Base-pair-resolution CRISPR-Cas technologies are producing a rapidly growing toolbox to induce precise genetic modifications^4^. Large-scale single cell CRISPR screening approaches in cell lines enable identification of key regulatory elements and their genes^5^. Extensions of these methods have been applied to primary human hematopoietic stem cells with base-pair resolution to directly confirm variant-function links^6^. However, depending on the targeted site and CRISPR-Cas machinery, editing efficiency can be highly variable which limits the power to detect functional effects. CRISPR screens might be more powerful if single cell data could be obtained that effectively discriminates between edited and non-edited cells.

Current methods assume high editing efficiency and achieve single-cell resolution by sequencing single guide RNA (sgRNA) as a proxy for genomic editing. However, these techniques cannot identify bystander and heterozygote editing thereby reducing power. They also rely on viral transduction which may confound cell-state-specific effects and remain challenging in primary cells^7^. An alternative strategy is to edit variants with non-viral methods such as electroporation or nanoparticle-mediated delivery of editor ribonucleoproteins or mRNA, but at the potential cost of lower efficiency^8^, and without the benefit of single cell resolution.

To address these challenges, we propose that defining the functional effects of CRISPR editing requires genomic DNA information at the single cell level in parallel with multimodal functional readouts. Doing this at scale is difficult since isolation of genomic DNA alongside mRNA is hindered by incompatible chemistries and reaction conditions. Previous iterations of DNA and RNA single cell approaches have relied on low-throughput and costly physical separation or targeted amplification methods^9,10^. Recent publications have increased throughput of whole genome and transcriptome sequencing techniques using transposase treatment of histone-depleted nuclei^11,12^. However, whole genome sequencing approaches cannot reliably recover information about specifically targeted loci.

We developed a plate-based method for the **M**ulti-omic **I**nvestigation of **N**ucleotide **E**diting with **CR**ISPR by **A**DT, **F**low Cytometry, and **T**ranscriptome sequencing (MINECRAFTseq, **Figure 1a)**. In each cell, it assesses successful editing by targeted genome sequencing **(Figure 1b)** and downstream functional effects in mRNA and cell surface protein phenotypes. It does not require any specialized equipment and can be easily implemented in any genomic editing approach. Alongside the experimental protocol, we demonstrate the power of single cell statistical models to define functional effects of individual non-coding alleles.

**Figure 1:**
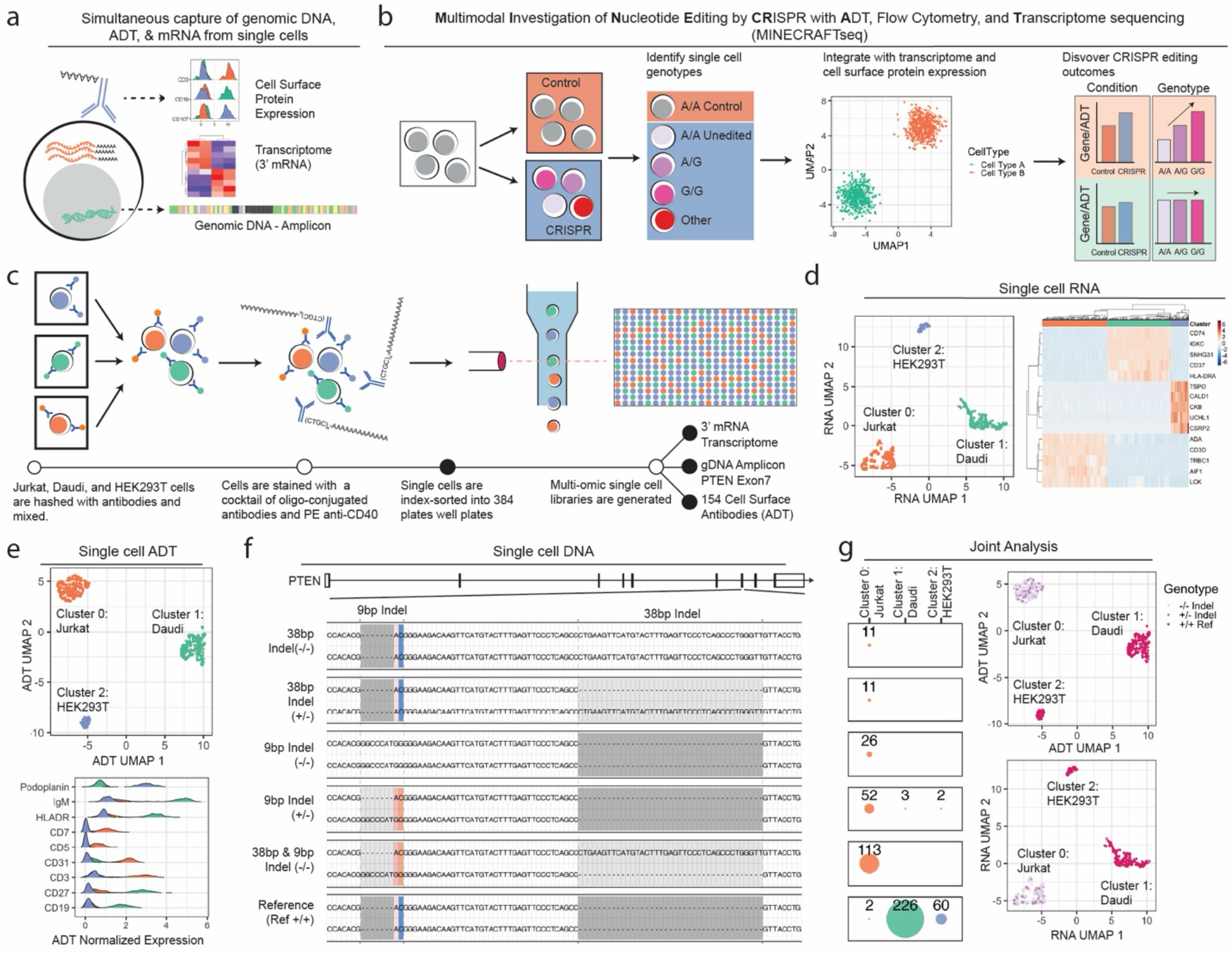
Simultaneous capture of single cell RNA, genomic DNA amplicons, and ADT with MINECRAFTseq. (a) MINECRAFTseq captures targeted genomic DNA amplicons, 3’ mRNA, and cell surface protein expression by ADT. (b) Deconvolution of CRISPR editing is used to identify genotypes and outcomes in mRNA and ADT expression. (c) Schematic overview of experimental design with a mixture of cell lines. (d) UMAP representation of the top 3000 variable genes following PCA, harmonization by plate, and clustering with SNN. Heatmap of differential gene expression between clusters. Scaled and log(CP10K +1) normalized values are plotted. Colors represent UMAP defined clusters (green = Daudi, red = Jurkat, blue = HEK293T). (e) UMAP representation of 154 ADTs following PCA, harmonization by plate, and clustering with SNN. Histograms of CLR normalized ADT values per cluster are plotted. (f) Identified unique PTEN genotypes from single cells. An amplicon flanking exon 7 of PTEN was used to identify genotypes. Two unique insertion/deletions (indel) are present. (g) The number of cells recovered per genotype is shown faceted by ADT cluster representing cell type. UMAP of ADT and mRNA embeddings with overlayed genotype. In UMAP representations each point represents a single cell.

### A robust multimodal single cell approach

MINECRAFTseq is a multimodal single cell assay that sequences genomic DNA amplicons, whole transcriptome RNA, and oligonucleotides tagging surface marker antibodies (Antibody-derived tags, ADTs) alongside flow cytometry-based cell-hashing (**Figure 1a**, **Methods**). It is efficient, allowing for the processing of thousands of cells per week at a cost of approximately ∼$3 per cell. MINECRAFTseq utilizes a modified FLASHseq full-length RNA sequencing protocol to yield high quality single cell transcriptomes^13^. Key modifications include the use of a nested PCR reaction to amplify the genomic DNA region flanking the target site, 3’ mRNA sequencing with barcoded oligoDT, and recovery of 154 oligo-conjugated antibodies for cell surface protein expression analysis (**Extended Data Figure 1**). This plate-based approach uses indexing flow cytometry to hash information on multiple conditions, cell types, and individuals within the same plate (**Figure 1c**), which allows us to control for plate effects and other important cell-level covariates.

We applied MINECRAFT-seq to a pool of unedited Jurkat, Daudi and HEK293T cells sorted into two 384-well plates **(Figure 1c** & **Extended Data Figure 2**). Previous reports have identified two insertion/deletion (indel) events in exon 7 of *PTEN* in Jurkat cells^14^. For this analysis, we amplified a genomic amplicon in *PTEN* exon 7 alongside whole transcriptome and protein expression data from a panel of 154 antibodies. Across both plates, an average of 58% of RNA reads aligned to the transcriptome. 555 out of 768 cells passed stringent RNA quality control and we recovered a mean of 5089 (+/-71 SEM) genes and 57540 (+/-2034 SEM) UMIs per cell, which is comparable to data from the 10x Genomics platform **(Methods, Extended Data Figure 2)**. Clustering these 555 cells on the 3000 most variable genes, we identified three populations with gene markers reflecting cell-line identity, for example *IGKC* (Daudi), *CD3D* (Jurkat), and *CKB* (HEK293T) (**Figure 1d**).

Of the ADT reads sequenced across both plates, 46% aligned to the ADT reference. Clustering the same 555 cells by ADT markers identified three populations with markers for B cells (Daudi, IgM+CD19+HLADR+), T cells (Jurkat, CD3+CD5+CD7+), and endothelial cells (HEK293T, Podoplanin+) (**Figure 1e**). We also show that CD40 ADT read counts correlate with flow cytometry-based antibody staining (R = 0.59) and gene expression (R = 0.36) (**Extended Data Figure 2)**. Overall, RNA, ADT, and flow cytometry modalities lead to almost perfectly concordant cell type calls (**Extended Data Figure 2**).

We obtained high-quality targeted DNA data. We called *PTEN* genotypes in 629 of 768 cells by mapping an average of 810 DNA reads per cell to the amplified region **(Methods, Extended Data Figure 2**). From our single cell DNA data, we confidently call the two previously reported indels in *PTEN*: (1) a 7bp insertion and 2bp mutation and (2) a 38bp insertion **(Figure 1f)**. As expected, we identify *PTEN* mutations in Jurkat and not Daudi or HEK293T cells **(Figure 1g** & **Extended Data Figure 2**). We detected miscalled genotypes in Daudi (1.3%) and HEK293T cells (3.3%) at very low rates, indicative of our platform’s high DNA calling accuracy **(Figure 1g).**

Surprisingly, our experiment revealed single cell heterogeneity in Jurkat genotypes. To confirm this observation, we sequence *PTEN* amplicons in bulk Jurkat cells. All three alleles representing reference, 9bp indel, and 38bp indel were detected at allele frequencies identical to the single cell data (**Extended Data Figure 3**). We observed no association between genotype and *PTEN* gene expression (**Extended Data Figure 2**).

### Identifying genotype-dependent outcomes

Next, we tested the power of MINECRAFTseq to resolve genome editing outcomes. In CRISPR editing experiments, cells are cultured before analysis and the secreted factors induced by successfully editing may alter the phenotype of neighboring cells. To resolve these differences, MINECRAFTseq can identify changes that are linked to genotype and not the broader cell culture environment. This analysis is performed by comparing the CRISPR condition, regardless of genotype, to the non-targeting control. These results are then compared with genotype-dependent changes identified only within the CRISPR condition (**Figure 1**). With this framework, we examined two distinct scenarios: 1) editing a regulatory region upstream of *HLA-DQB1* and assessing genome-wide expression effects and 2) disrupting *FBXO11* and assessing response after CD40L stimulation.

We previously identified that a variant, rs71542466, in *HLA-DQB1* regulated gene expression in a context-specific manner^15^. Here, we targeted this variant using CRISPR-Cas mediated homology-directed repair (HDR) in two cell lines, HH and Daudi, with ribonuclear proteins and repair template **(Figure 2a** & **Extended Data Figure 4)**. We stain cells with a fluorophore-conjugated anti-HLA-DQ antibody since we did not include it in our pre-selected ADT panel. Additionally, we used fluorophore-conjugated antibodies to tag non-targeted cells to detect them with indexing flow cytometry (**Extended Data Figure 4**). We label cells before mixing and staining with the ADT antibody panel to ensure consistent surface marker staining.

**Figure 2:**
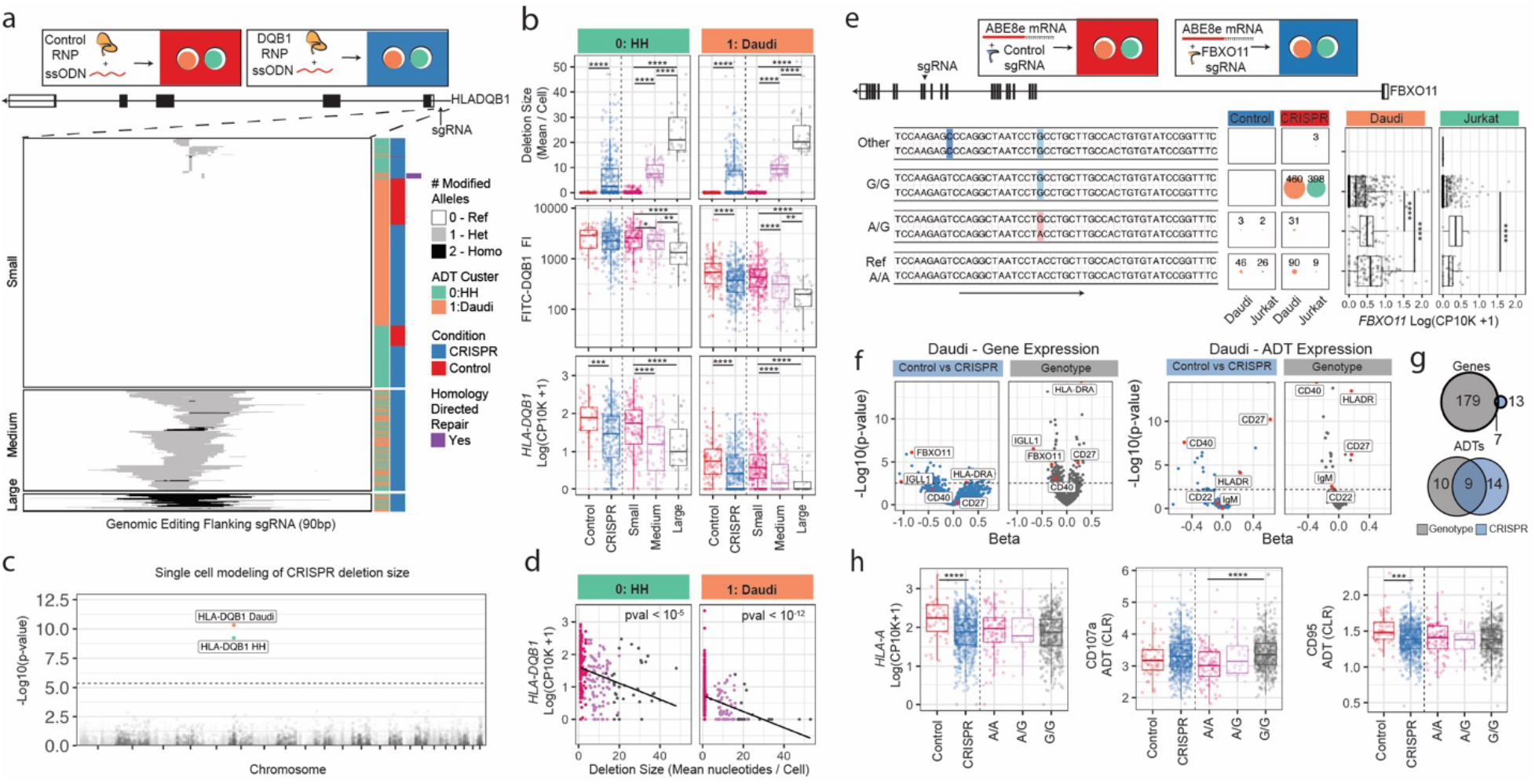
MINECRAFTseq identifies genotype dependent editing outcomes in single cells. (a) Schematic of genomic editing in the 5’UTR of HLA-DQB1 in HH and Daudi cells. CRISPR-Cas9 RNP along with HDR donor material was used to induce the rs71542466 (C to G) variant in HLADQB1. Heatmap of editing in the locus around the sgRNA. Each row represents an individual cell. Color represents heterozygote(grey) or homozygote(black) modification at that position. Conditions and cell types clustering is displayed (right) with editing clusters (left). (b) Mean deletion size, indexing cytometry FITC-DQB1 flourescence intensity(FI), and normalized *HLA-DQB1* gene expression is plotted per condition, cell type, and editing cluster. (c) Manhattan plot of p-values generated from single cell linear modeling of gene expression to log normalized deletion size in Daudi and HH cells. Dotted line represents the FDR cut-off of 0.05. Each point represents a gene. (d) Scatterplot of deletion size to normalized *HLA-DQB1* gene expression. (e) Schematic of FBXO11 CRISPR base editing in Jurkat and Daudi cells. ABE8e base editors were used to induce a A-G change causing an alteration in splicing. Genotypes (>2 cells) from the FBXO11 amplicon around the sgRNA (red arrow) are shown with number of cells per genotype (right) and normalized expression of *FBXO11* by genotype faceted by ADT cluster representing cell type. (f) Differential gene and ADT expression in Daudi cells identified with linear modeling of control vs CRISPR cells and in a dosage-dependent model from only CRISPR edited cells. Dotted line represents the FDR cut-off of 0.05. (g) Comparison of significant genes and ADTs between control vs CRISPR and genotype-dependent analysis. (h) Normalized expression of *HLA-A*, CD107a, and CD95. P-values are derived from likelihood ratio statistics from linear modeling with values of * < 0.05, **<0.01, ***<0.001, & ****<0.0001.

We observed highly heterogeneous editing in both cell lines with DNA deletions of variable size **(Figure 2a&b**). We removed cells with low quality RNA profiles along with 15 cells with DNA insertions (**Extended Data Figure 4**). We used K-means clustering to group DNA edits into three classes: small, medium, and large **(Methods)**. After identifying cell types by clustering on ADT markers (**Extended Data Figure 4**), we observed a strong association between DNA deletion class and *HLA-DQB1* expression at the transcript (ANOVA p < 0.0001) and protein level (ANOVA p <0.0001) (**Figure 2b).**

We wanted to assess whether editing this regulatory region had a specific effect on *HLA-DQB1* or on other genes. We modeled the relationship between read counts and log2 deletion size in HH and Daudi cells in 4367 and 3621 genes passing quality control respectively, using a negative binomial model **(Methods)**. Strikingly, we observed that deletion size is specifically associated with only *HLA-DQB1* counts(p<10^-8^) (**Figure 2c&d**) and no other gene or protein (FDR > 0.05). These results demonstrate the specific genomic function that can be identified with MINECRAFTseq. Finally, we observed HDR-edited cells with the intended C to G transition in 10 of 646 cells **(Figure 2a)**. Given this low efficiency, we did not detect statistically significant changes in gene or protein expression (**Extended Data Figure 4**).

Next, we explore the ability to detect stimulation-induced transcriptome-wide changes after disrupting *FBXO11*, an intracellular signaling protein. We used ABE8e base editors to induce an A to G transition in two cell lines to introduce a loss of function allele **(Figure 2e** & **Extended Data Figure 5**). We identified four genotypes corresponding to editing at the targeted nucleotide **(Figure 2e)**. We observed homozygous editing at the target nucleotide in 86% of the cells. We removed cells with bystander editing outcomes for downstream analysis. At the single cell level, we confirm that disruption of splicing impacts gene abundance with a genotype-dependent 2.2-fold decrease in *FBXO11* transcripts in G/G vs A/A Daudi cells **(Figure 2e)**.

We expected that the introduction of the allele and subsequent loss of *FBXO11* mRNA would blunt the response to CD40L stimulation. To identify these changes, we first used a negative binomial single-cell model to compare the expression of 3427 genes passing quality control thresholds and 154 ADT surface markers in control vs CRISPR cells after stimulation (**Methods**). In this genotype-unaware analysis, we discover 20 significant genes (FDR < 0.05) and 23 significant ADT protein tags (FDR <0.05) when controlling for plate and library complexity **(Figure 2f)**. However, our approach allows identification of specific genotype-dependent effects. In this approach, we examine cells only in the CRISPR condition. To increase power, we model allele dosage – that is 0 for reference, 1 for heterozygotes, or 2 for homozygote edited. Importantly, modeling genotype-specific effects idenfied broader functional effects, revealing 186 genes (FDR<0.05) and 19 proteins (FDR<0.05) whose expressino varied with edit dosage **(Figure 2g)**. With this modeling, we observed significant down regulation of *FBXO11, IGLL1*, CD40, and IgM **(Figure 2f)**, indicating the expected overall decrease in CD40L-mediated activation with negative feedback on CD40 gene and protein expression.

Surprisingly, we identify a subset of genes and proteins that are downregulated in the CRISPR condition reflecting non-specific environmental effects that have no genotype dependence such as *HLA-A* (Likelihood Ratio Test (LRT)-culture-condition p-value < 10^-4^) and CD95 (LRT culture-condition p-value < 10^-5^) **(Figure 2h)**. FBXO11 has been described as a regulator of CD95 protein expression^16^, but our results suggest that this may be an indirect effect. In contrast, we observed genotype-dependent effects obscured in condition-specific analysis such as an increase in CD107a protein expression (LRT genotype p-value < 10^-4^) **(Figure 2h).**

### Knockout in primary human CD4 T cells

We assessed whether MINECRAFTseq can be used to investigate functional variants in primary human cells. We focused specifically on T cells and *PTPRC*, the gene encoding CD45, since it is linked with inflammatory disorders including asthma, Crohn’s disease, and rheumatoid arthritis^17–20^. To study the role of *PTPRC* in CD4 T cell activation, we isolated naive CD4 T cells from three individuals and introduced a premature stop codon with BE4 base editors **(Figure 3a)**. We sorted four 384-well plates from three individuals, with 628 cells passing QC. We used fluorophore conjugated antibodies to track editing versus control conditions and individuals; we also confirmed assignment of individuals using genotype data derived from sequencing data **(Methods. Extended Data Figure 6**)

**Figure 3:**
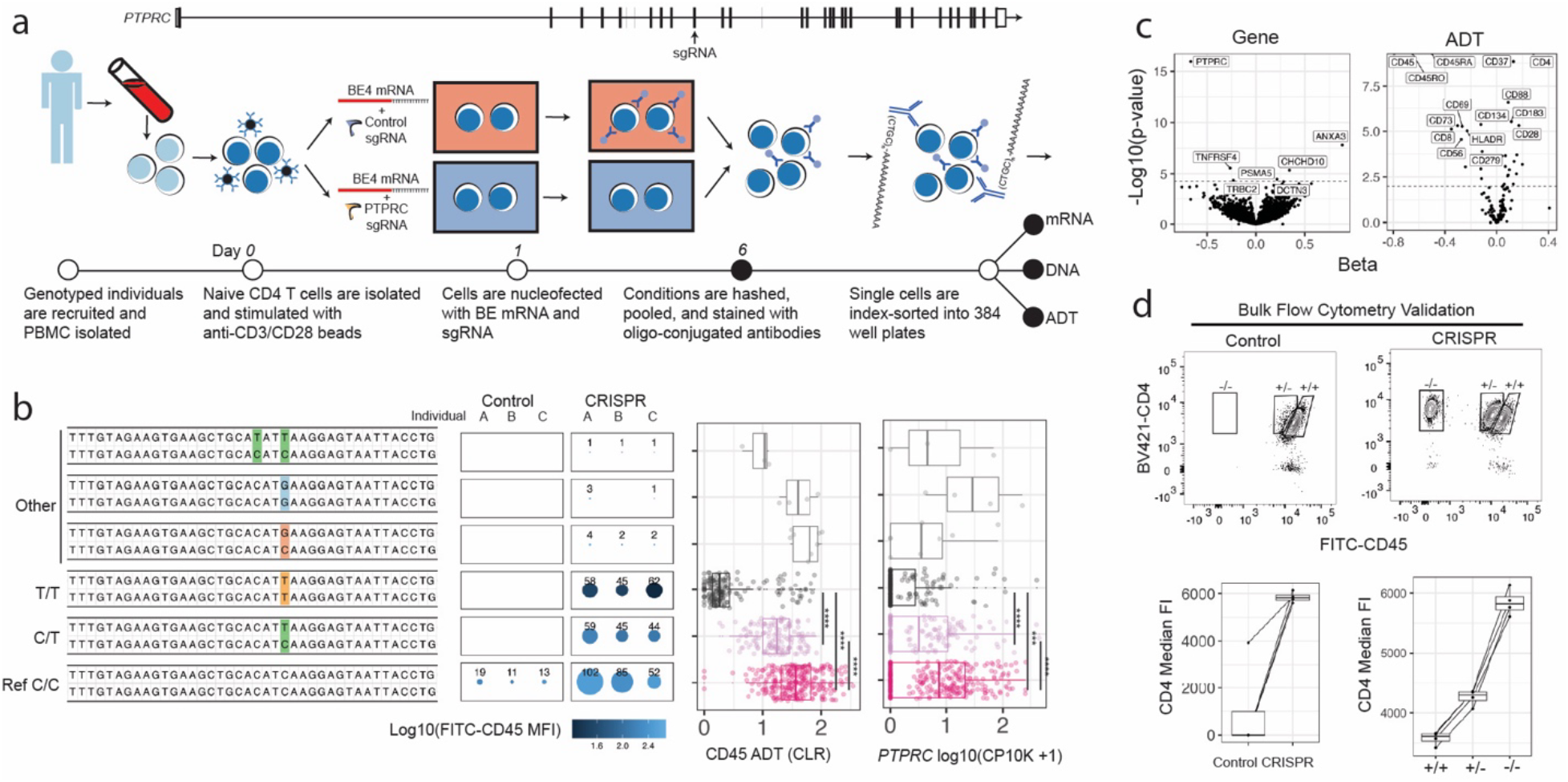
Induction of a premature stop codon in *PTPRC* identifies changes in gene and cell surface protein expression. (a) Schematic of genomic editing in the *PTPRC* in primary human naïve CD4 T cells. BE4-NG base editors were used to induce a premature stop codon. (b) Genotypes (>2 cells) from the PTPRC amplicon around the sgRNA are shown with number of cells per condition and individual (right) and normalized expression of *PTRPC* by genotype. (c) Volcano plot of differential gene expression and ADT expression identified with linear modeling. P-values and beta from likelihood ratio tests are plotted for the genotype term. Dotted line represents the FDR cut-off of 0.05. (d) Confirmation of results in bulk edited CD4 T cells with flow cytometry. Representative plots of CD4 vs CD45 is shown for control and CRISPR conditions. The combination of CD4 and CD45 cell surface expression is used to identify heterozygote and homozygote edited cells (+/-, -/-). P-values are derived from likelihood ratio statistics from linear modeling with values of * < 0.05, **<0.01, ***<0.001, & ****<0.0001.

We observed that the T allele dramatically abrogates CD45 protein expression in an allele-dependent manner (**Figure 3b**). Transcript abundance of *PTPRC* is reduced, likely due to nonsense mediated decay^21^ (**Figure 3b**) (LRT p-value < 10^-15^ beta = -0.66). Given the large effect on *PTPRC* gene and CD45 protein expression and the central role they play in T cell biology, we predicted that this loss of function allele would induce *trans* effects. Hence, we tested 5544 genes and 154 proteins for genotype-dependent (dosage) effects (**Methods**).

Disruption of *PTPRC* altered 32 cell surface proteins (FDR<0.05) including increased expression of activation markers such as CD25 and CD28 (**Figure 3d**). Indeed, when we projected the cells into a uniform manifold approximation and projection (UMAP) embedding based on ADT expression, edited cells clustered separately (**Extended Data Figure 7&8**). The *PTPRC* loss of function allele also had broad effects on gene expression, with 60 genes demonstrating differential effects (FDR<0.2). GSEA analysis of these genes identifies an enrichment for Th1 and activation pathways (**Extended Data Figure 9**)

As a key example of a trans effect, we observed that loss of function allele in *PTPRC* dramatically reduced the gene*TNFRSF4* located 18Mb from *PTRPC* (LRT p-value < 10^-5^ beta = -0.27), which is a critical T cell co-stimulatory molecule expressed on Tregs^22^. Consistent with this observation, we also saw that *PTPRC* loss of function reduces surface marker expression for CD134(OX40), encoded by the *TNFRSF4* gene (LRT p-value < 10^-5^, beta = -0.12). (**Extended Data Figure 10**).

We observed that PTPRC also induced CD4 protein expression (LRT p-value < 10^-16^, beta = 0.28). To validate this intriguing observation, we examine CD4 cell surface expression in bulk CRISPR-edited CD4 T cells (**Figure 3e**). We confirm editing efficiency by flow cytometry to assay CD4 and CD45 protein expression alongside amplicon sequencing (**Extended Data Figure 11**). Consistent with the single cell results, we observe 1.5-fold increase in CD4 cell surface expression between control and CRISPR conditions (p-value = 0.017, Wilcoxon Rank Test). Interestingly, analysis of co-expression of CD4 and CD45 from genomically edited cells allowed for the identification of a heterozygous-edited population by flow cytometry. For the first time, we show that loss of *PTPRC* in naïve CD4 T cells leads to an upregulation of CD4 protein expression, consistent with previous reports that CD4 and CD45 co-localize on the cell surface to regulate T cell activation^23,24^.

As before, we assessed condition-specific (control vs CRISPR) effects (**Extended Data Figure 12**). We observed broad changes in IFN response genes (*IFI6*, *ISG20, ISG15,* LRT FDR<0.05). These genes likely reflect non-specific changes caused by culture conditions that would confound bulk editing assays. In contrast, we observe no change in *PTPRC* (FDR > 0.05).

### Defining the function a causal allele in the RPL8 locus

After confirming the effects of a loss of function allele, we wanted to genomically modify a variant that controls gene expression. We identified rs2954658 (T to C transition) within the *RPL8* locus that is associated with increased expression in previous B cell eQTL studies; this allele was suggested to be the likely causal allele by statistical finemapping analyses^25,26^ (**Extended Data Figure 13**). To assess if the allele has a true functional effect, we edited Daudi cells from T to C at this single base site. We sorted single cells into 6 384-well plates, applied MINECRAFT-seq and analyzed 888 cells passing quality control. Using single cell genotype information, we confirmed a strong statistical effect of the T allele on *RPL8* expression (p<10^-6^, 1.1-fold-change, **Figure 4a**). We tested 4106 genes and 154 proteins for a genotype effect; we did not observe a genotype effect on any other individual gene (FDR > 0.05), again highlighting the ability of our approach to identify a specific effect (**Figure 4b**). We identified no other with culture-condition analysis (FDR>0.05).

**Figure 4:**
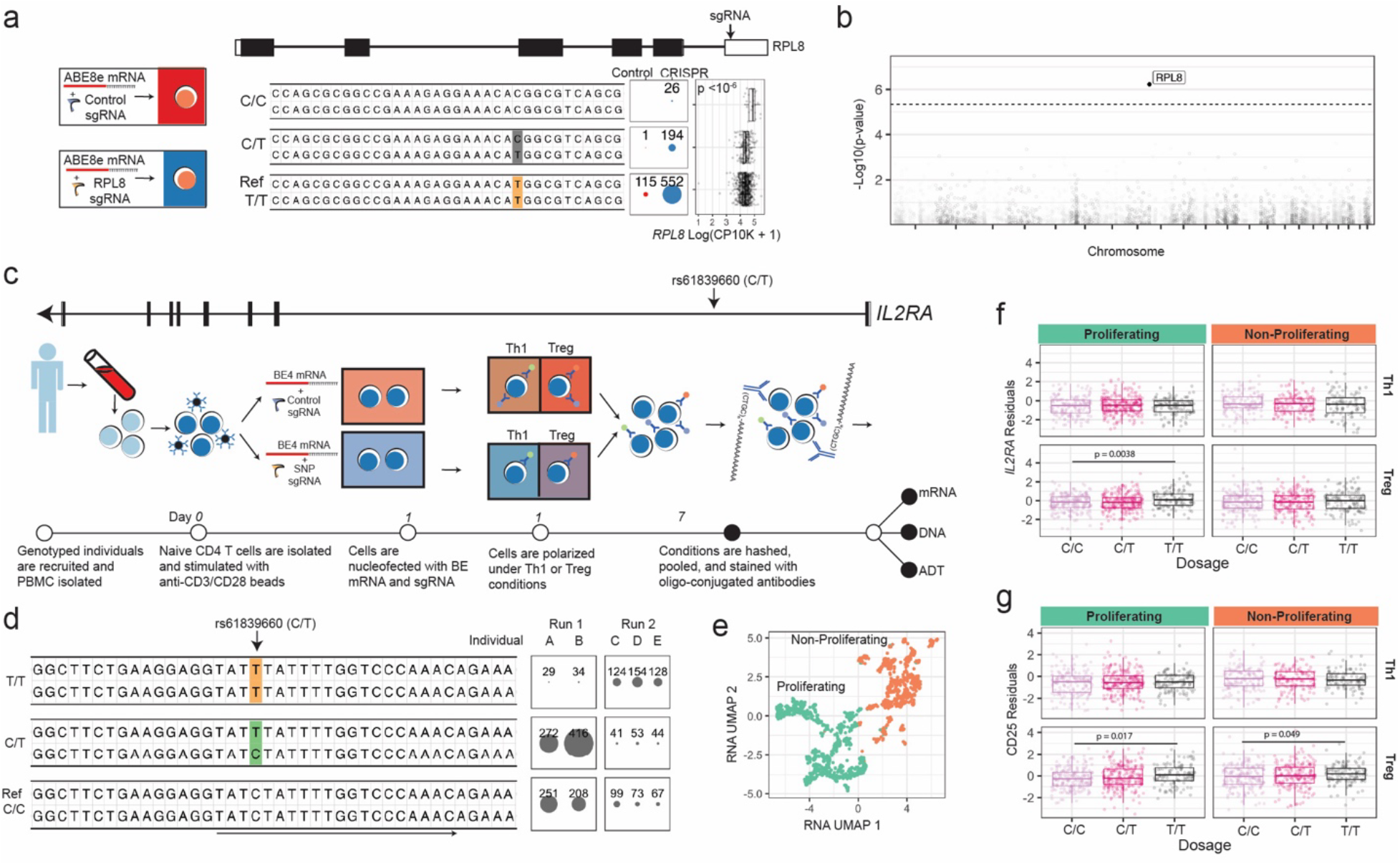
Identification of cell-state specific effects variant effects with MINECRAFTseq. (a) Schematic of genomic editing in the RPL8 in Daudi cells. ABE8e base editors were used to induce the rs2954658 (T alternative to C reference) variant. Genotypes (>2 cells) from the RPL8 amplicon around the sgRNA are shown with number of cells per condition (right) and normalized expression of *RPL8* by genotype. (b) Manhattan plot of p-values generated from single cell linear modeling of gene expression to dosage at the targeted nucleotide. Dotted line represents the FDR cut-off of 0.05. Each point represents a gene. (c) Schematic of genomic editing of rs61839660 in primary human naïve CD4 T cells polarized to Th1 or Treg phenotypes. BE4-NG base editors were used to induce the alternative T allele in genotyped non-autoimmune individuals. (d) The three major genotypes from the amplicon around the sgRNA are shown with number of cells per run and individual (right) and normalized expression. (e) UMAP of top 3000 variable genes scaled, normalized, and harmonized by plate identifies two clusters. (f) Residuals of *IL2RA* gene expression with plate, individual, and library size regressed. (g) Residuals of CD25 ADT expression with plate, individual, and library size regressed. P-values from likelihood ratio tests are plotted. Th1 and Treg polarizations were identified with indexing flow cytometry. P-values from likelihood ratio tests are plotted.

As reported previously^27^, Daudi cells encompass a heterogenous population (**Extended Data Figure 14**). Clustering on ADT expression identifies two cell states. Modeling *RPL8* gene expression in each cluster revealed no differences in association to genotype in the two cell states (LRT interaction p=0.3), **Extended Data Figure 14**).

### A cell-state specific IL2RA regulatory variant

Finally, we examine the effects of a non-coding autoimmune variant in *IL2RA*, rs61839660, associated with type 1 diabetes, asthma, systemic lupus erythematosus, and rheumatoid arthritis^19,28–30^. Previous studies have reported that this variant may be causal and linked to downregulation of *IL2RA* gene expression and Treg differentiation^31^. However, others have shown that the haplotype associated with this variant increases IL2RA protein and gene expression^32,33^. Given these contradictory findings, we hypothesize that rs61839660 regulates *IL2RA* and CD25 expression in a highly context-dependent manner. Hence, we investigated the effect of this variant in primary T cells polarized towards Th1 or Treg states **(Figure 4c)**.

We recruited five individuals and use BE4 base editor to induce a C to T transition. We analyzed a total of 1993 cells across two experiments after QC and removing cells with bystander editing outcomes **(Figure 4d).** After harmonizing by experiment and plate, we create a joint embedding from differentially expressed mRNA and identify three prominent clusters that separate on proliferative status (*MKI67* and *ACTB*) **(Figure 4e** & **Extended Data Figure 15)**.

Using a single cell model, we demonstrate that the T allele is associated with a significant increase in *IL2RA* only in proliferating Treg-polarized cells (LRT p-value = 0.0038, 1.12-fold change, **Figure 4f**). The allele also effects expression of CD25, encoded by *IL2RA*, in non-proliferating (LRT p-value = 0.0175, beta = 0.10) and proliferating (LRT p-value = 0.0498, beta = 0.0917) Tregs. However, genotype has no effect in Th1-polarized cells in either CD25 protein or *IL2RA* (LRT p-value >0.05, **Figure 4g**), indicating that this is a context-dependent regulatory variant.

Overall, we define a functional effect of the rs61839660 T risk allele in primary human CD4 T cells in a highly context-dependent manner and highlight the utility of MINECRAFTseq to discover the most subtle cell state specific changes caused by non-coding variants. The identification of this effect argues that the disease association of this locus is driven at least in part by this allele.

### Conclusions

Scalable and flexible methods are urgently needed to investigate the rapidly emerging list of genomic variants associated with human disorders. Functionalizing alleles is essential to furthering our understanding of the root genetic causes of disease. We present a highly customizable multi-omic approach applied to a multitude of genomic editing techniques in cell lines and primary immune cells. To our knowledge, this is the only technique capable of capturing genomic DNA amplicons alongside the transcriptome and cell surface protein expression at scale, providing invaluable multimodal readouts on cell state.

We envision that MINECRAFT-seq can be applied on a case-by-case basis to resolve any genetic editing outcome. These include the study of regulatory regions, gene disruption in intracellular and cell surface proteins, and base-pair level editing. We assert that it can define both the *cis* and *trans* functions of complex trait alleles in primary human cells. Our experiments demonstrate the statistical power advantage of single cell methods over bulk analysis, and the ability to localize highly specific effects to a single downstream gene.

**Extended Data Figure 1:**
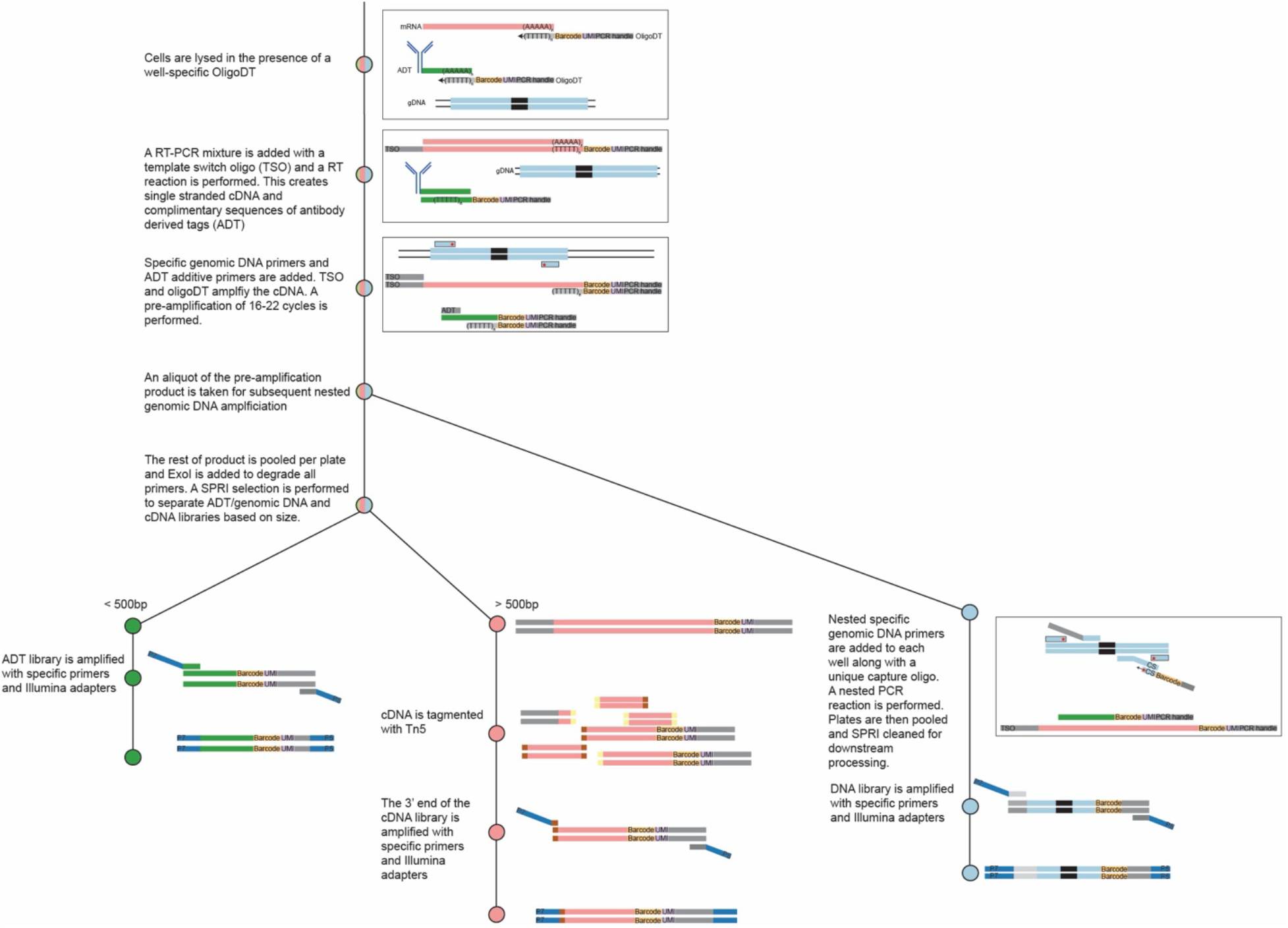
Schematic of library generation.

**Extended Data Figure 2:**
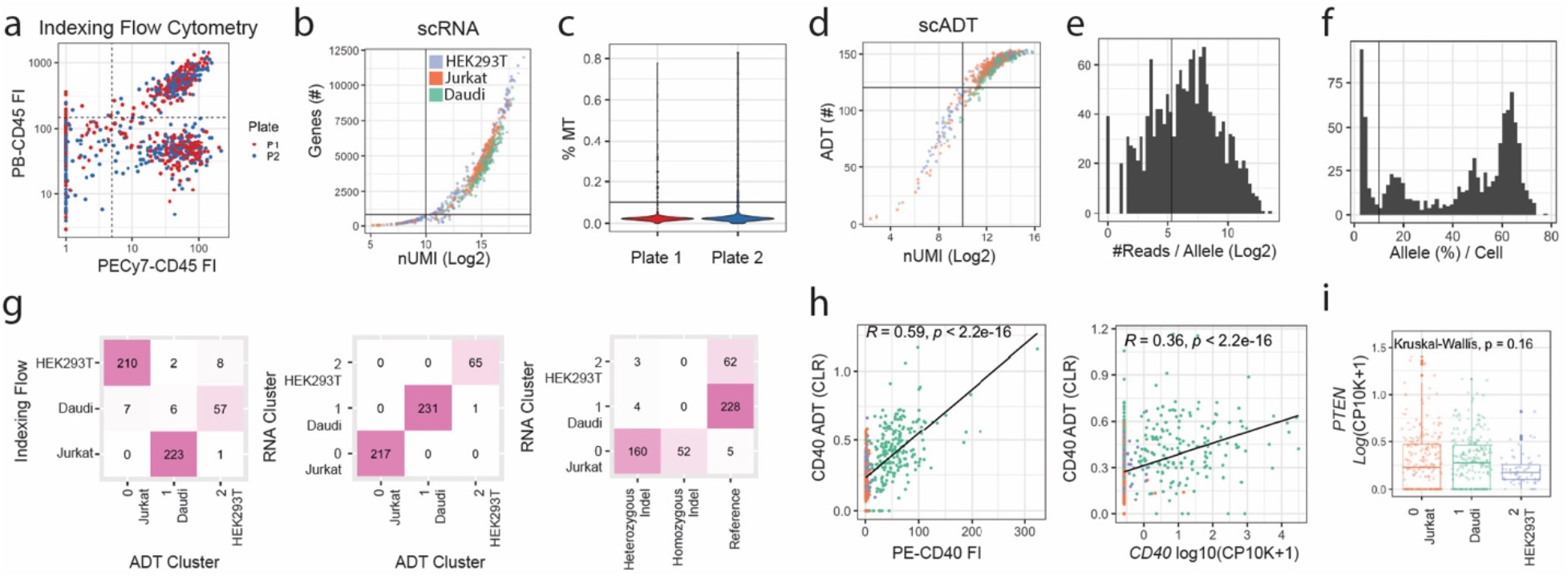
Indexing flow cytometry gating and quality control from two 384 well plates of mixed cell lines. (a) Indexing flow cytometry data is plotted with gating of each population shown (HEK293T PECy7-PB-; Jurkat PECy7+PB-; Daudi PECy7+PB+). (b) Single cell RNA metrics from both plates. Cells are colored by cell identity as gated in a. Lines represent the 1000 uMI and 1000 gene cutoff used to filter cells in downstream analysis. (c) Percent mitochondrial mRNA UMI per cell per plate. Horizontal line indicates the 10% filtering cutoff. (d) Single cell ADT metrics from both plates. Cells are colored by cell identity as gated in a. Lines represent the 1000 uMI and 120 ADT cutoff used to filter cells in downstream analysis. (e) Histogram of the number of PTEN aligned reads in the top two alleles from all cells. The horizontal line indicates the 40 read filtering minimum for removing low quality alleles. (f) Histogram of the percentage of alleles per cell in all cells. The horizontal line indicates the 10% allele cutoff. (g) Concordance plots of cell types defined by the different modalities. Color represents total cell number. (h) Correlation plots of CD40 protein expression by ADT sequencing, indexing flow cytometry and gene expression. R values and pearson correlations are displayed. Color represents cell type as defined by ADT cluster. Normalization approaches described in methods. (i) Log(CP10K +1) gene expression of *PTEN* per ADT cluster. Each point represents a single cell. P value of nonparametric Kruskal-Wallis test is displayed.

**Extended Data Figure 3:**
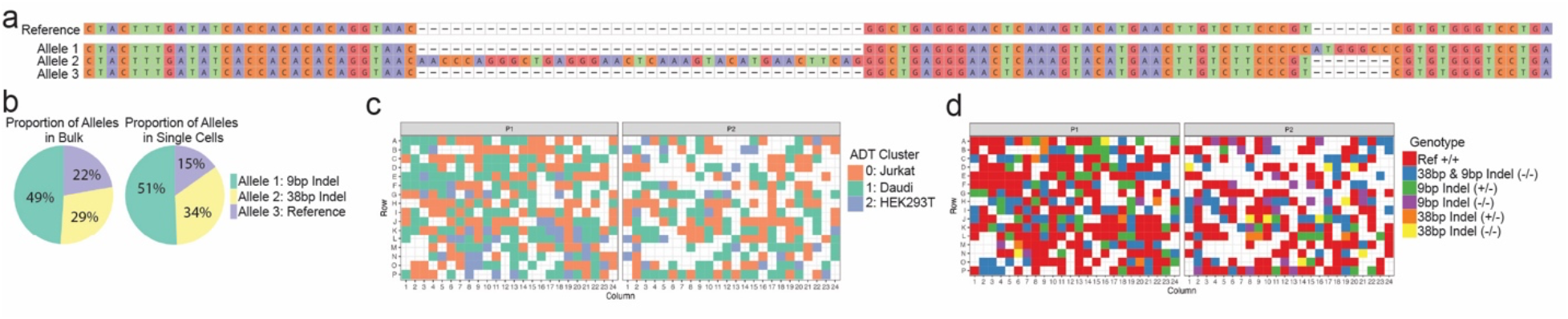
Bulk DNA sequencing of PTEN amplicons in Jurkats matches single cell readouts. (a) Representation of top three alleles recovered from bulk sequencing of PTEN amplicons from Jurkat cells. (b) Frequency of alleles in bulk data and single cell data. For single cell data, allele frequencies represent the sum of the identified allele in all cells divided by the total number of cells multiplied by 2. (c) Location of cell types as defined by ADT clusters across both plates. (d) Location of genotypes as defined by single cell gDNA sequencing of the PTEN amplicon.

**Extended Data Figure 4:**
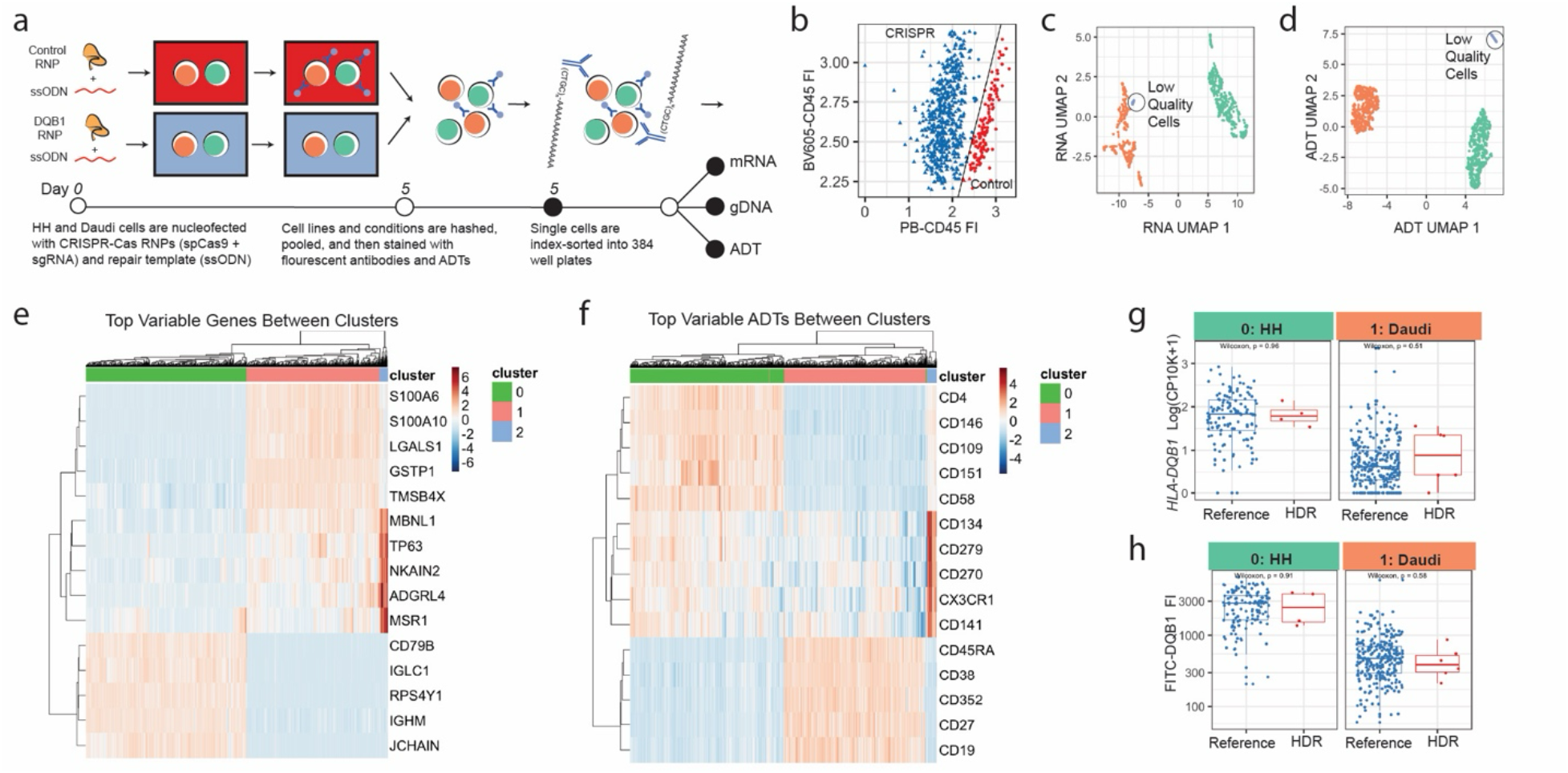
HDR corrected cells can be identified but are not different from controls. (a) Full experimental outline of HLADQB1 editing in HH and Daudi cells. (b) Representative plots of CRISPR vs control gating based on indexing flow cytometry. (c) UMAP of the top 3000 variable genes scaled, normalized, and harmonized by plate identifies three clusters (d) UMAP of all ADTs scaled, normalized, and harmonized by plate identifies three clusters. Low quality cells are labelled. (e) Heatmap of top variable genes, scaled and normalized within each cluster. (f) Heatmap of top variable ADTs scaled and normalized in each cluster. (g) Normalized *HLA-DQB1* gene expression in HH and Daudi cells between reference and HDR genotypes. (h) FITC DQB1 MFI in HH and Daudi cells between reference and HDR genotypes. Each dot represents a single cell. HDR corrected cells were all heterozygotes with one reference and one correct allele identified.

**Extended Data Figure 5:**
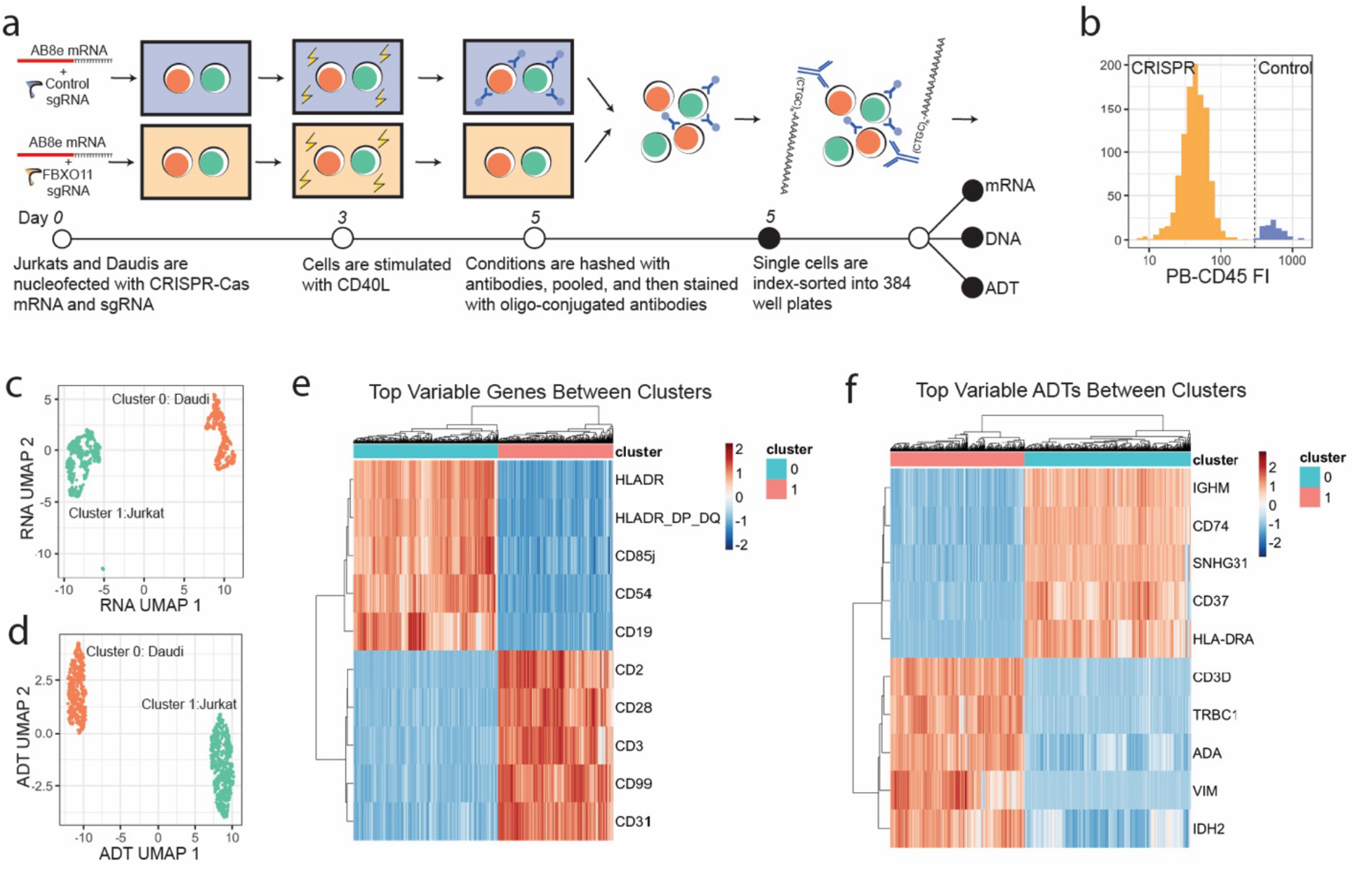
Indexing cytometry gating and cluster analysis of FBXO11 edited cells identifies two cell types separated by canonical markers. (a) Full experimental outline of FBXO11 editing in Jurkat and Daudi cells. (b) Representative plots of CRISPR vs control gating based on indexing flow cytometry. (c) UMAP of the top 3000 variable genes scaled, normalized, and harmonized by plate identifies three clusters. (d) UMAP of all ADTs scaled, normalized, and harmonized by plate identifies three clusters. (e) Heatmap of top variable genes, scaled and normalized within each cluster. (f) Heatmap of top variable ADTs scaled and normalized in each cluster.

**Extended Data Figure 6:**
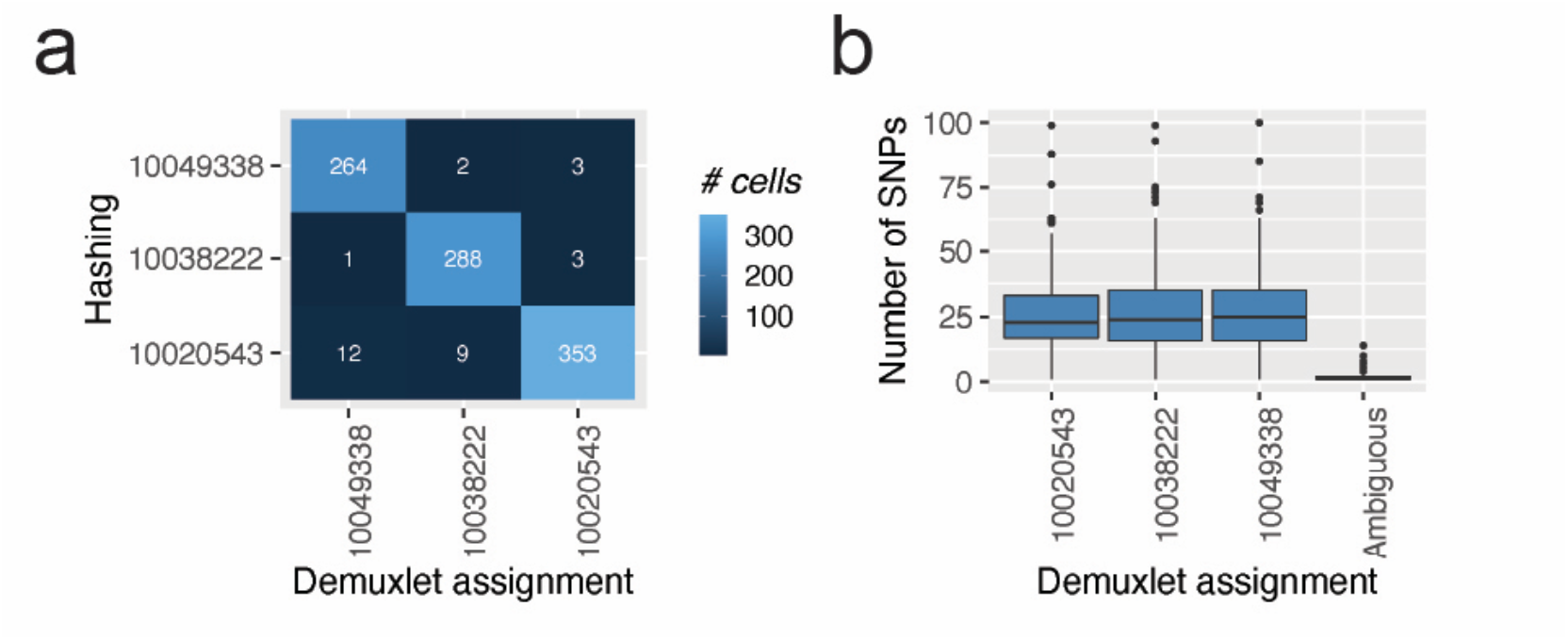
Index sorting based hashing is concordant with genotype-based demultiplexing from single cell RNAseq data. (a) Concordance of genotypes and hashing identify of the three individuals from Figure 4. (b) Number of SNPs identified in RNA sequencing data in each cell per individual.

**Extended Data Figure 7:**
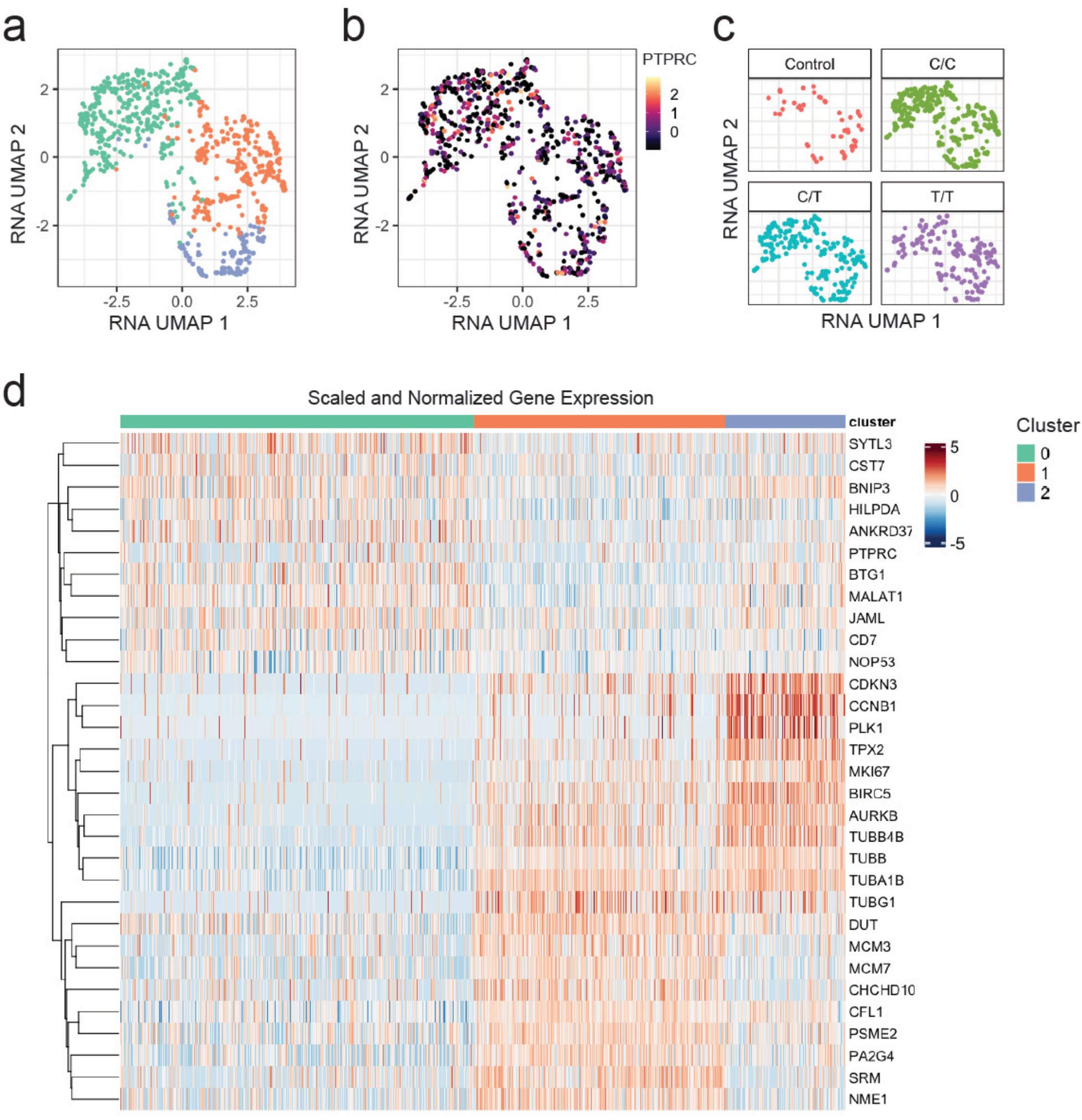
Clustering of *PTRRC* edited cells identifies broad changes in gene expression linked to proliferation and survival. (a) UMAP of the top 3000 variable genes scaled, normalized, and harmonized by plate identifies three clusters. (b) Scaled and normalized gene expression of *PTPRC*. (c) Distribution of cells based on condition and genotype in RNA UMAP embedding. (d) Heatmap of top variable genes, scaled and normalized within each cluster.

**Extended Data Figure 8:**
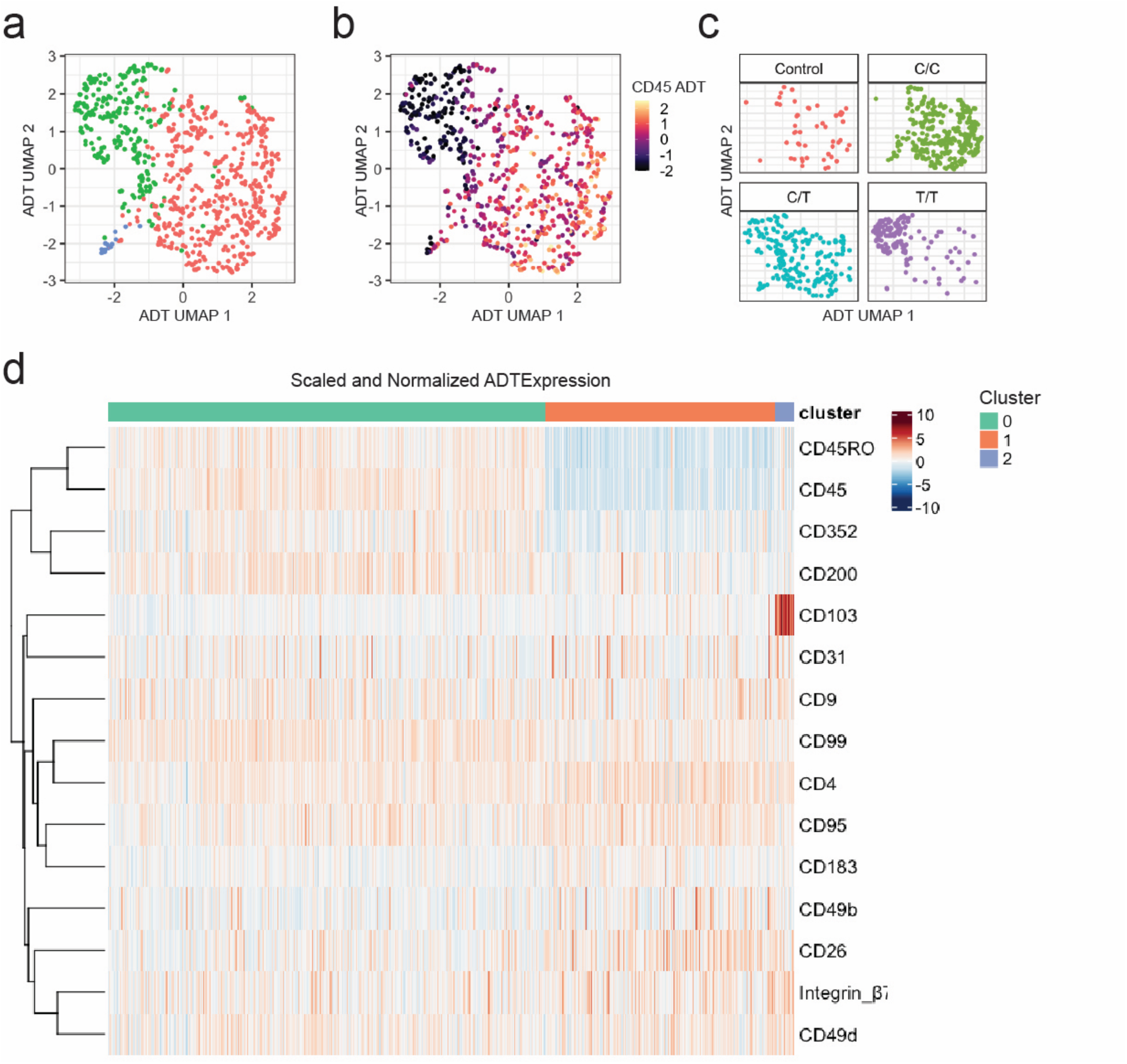
Analysis of *PTRPC* edited cells identifies a knockout cluster. (a) UMAP of all ADTs scaled, normalized, and harmonized by plate. (b) Scaled and normalized ADT expression of CD45. (c) Distribution of cells based on condition and genotype in ADT UMAP embedding. (d) Heatmap of top variable ADTs, scaled and normalized within each cluster.

**Extended Data Figure 9:**
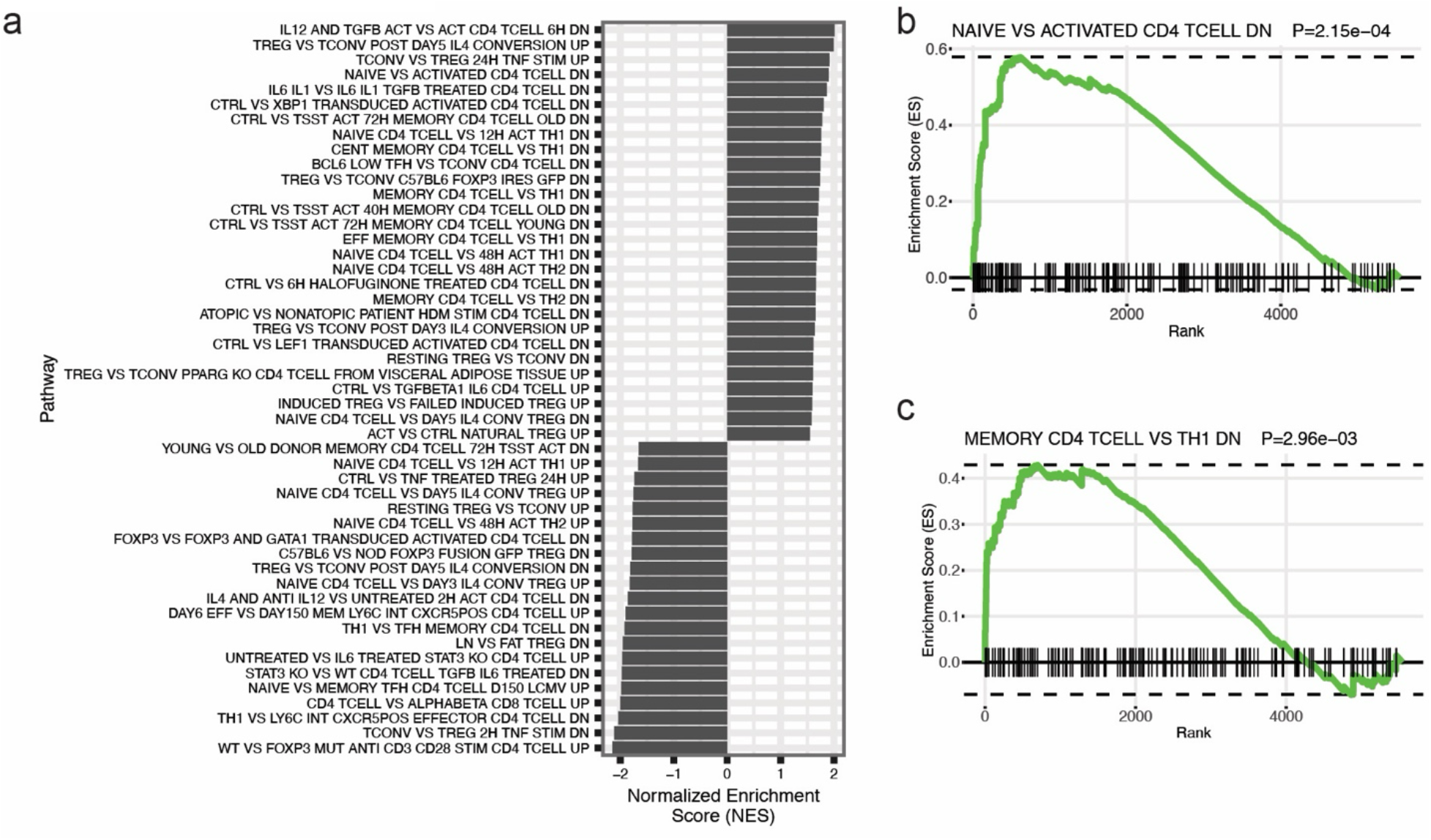
Gene set enrichment analysis identifies activation and Th1 programs in PTPRC knockout cells. (a) Normalized enrichment scores of significant G7 immunological pathways in CD4 T cells. P-values are derived from fgsea and are Bonferroni-corrected. (b) Enrichment plot of naïve vs activated CD4 T cells. (c) Enrichment pot of memory CD4 vs Th1 cells. P-values are derived from fgsea and are Bonferroni-corrected.

**Extended Data Figure 10:**
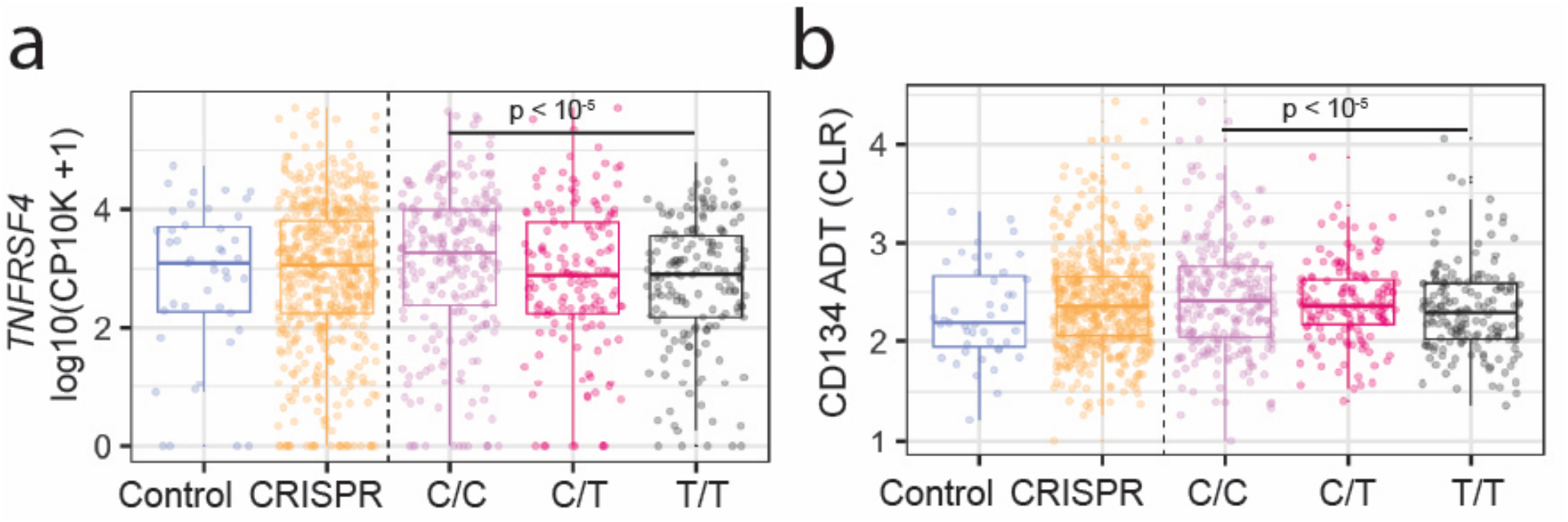
Expression of OX40 receptor is decreased in PTPRC edited cells. (a) Normalized expression of *TNFRSF4* by condition and genotype. (b) Normalized expression of CD134 by condition and genotype. P-values are derived from likelihood ratio statistics with linear modeling. No differences between control and CRISPR edited cells were detected (FDR > 0.05).

**Extended Data Figure 11:**
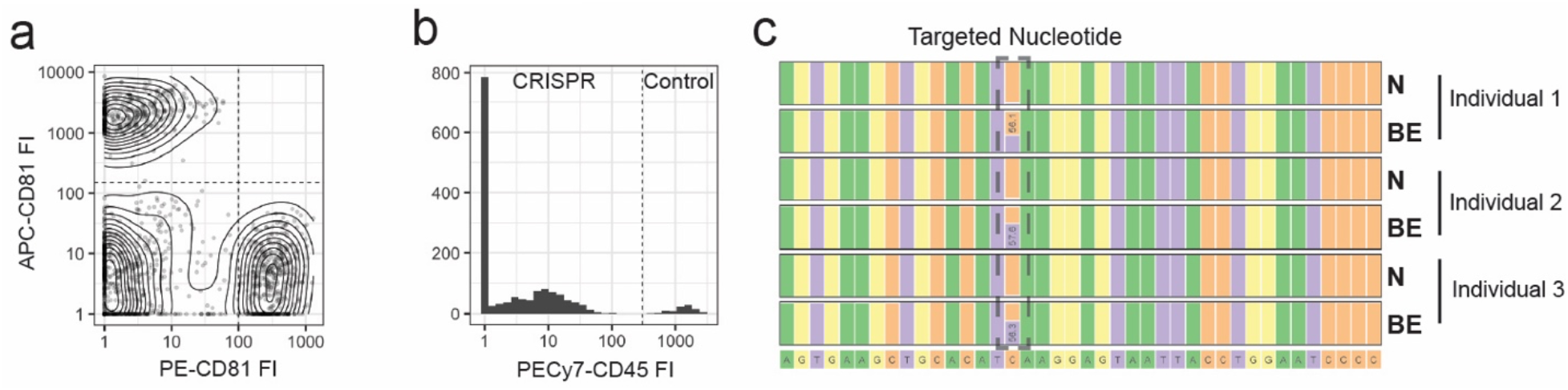
Indexing flow cytometry gating and bulk sequencing of CRISPR edits. (a) Representative plots of individual gating based on indexing flow cytometry staining with PE anti-CD81 and APC anti-CD81.Dotted lines represent the gates used to define each individual. (b) Representative plots of CRISPR vs control gating based on indexing flow cytometry of PECy7 anti-CD45. (c) Analysis of PTPRC amplicon from three bulk edited CD4 T cells. N represents non-targeting control and BE represents base-edited samples. A C to T conversion if noted in ∼ 50% of reads. Data was analyzed and plotted with CRISPResso2.

**Extended Data Figure 12:**
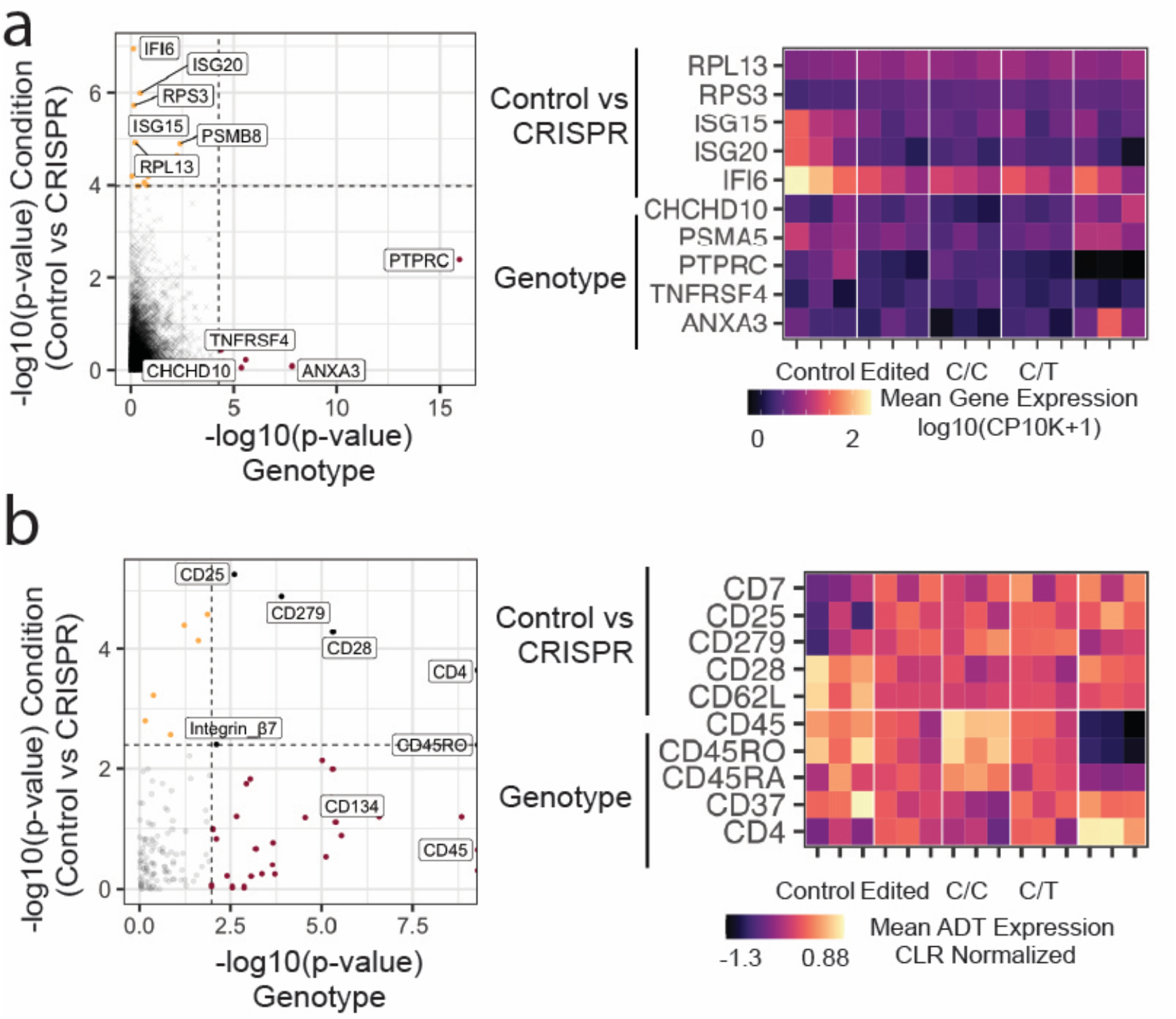
Differences between genotype and condition-specific analysis in PTPRC editing experiments. (a) Comparison of differential gene expression in control vs CRISPR and genotype-dependent analyses. Heatmap of normalized mean gene expression pseudobulked per individual and condition. (b) Comparison of ADT gene expression in control vs CRISPR and genotype-dependent analyses. Heatmap of normalized mean ADT expression pseudobulked per individual and condition. P-values from likelihood ratio tests are plotted. Dotted line represents the FDR cut-off of 0.05.

**Extended Data Figure 13:**
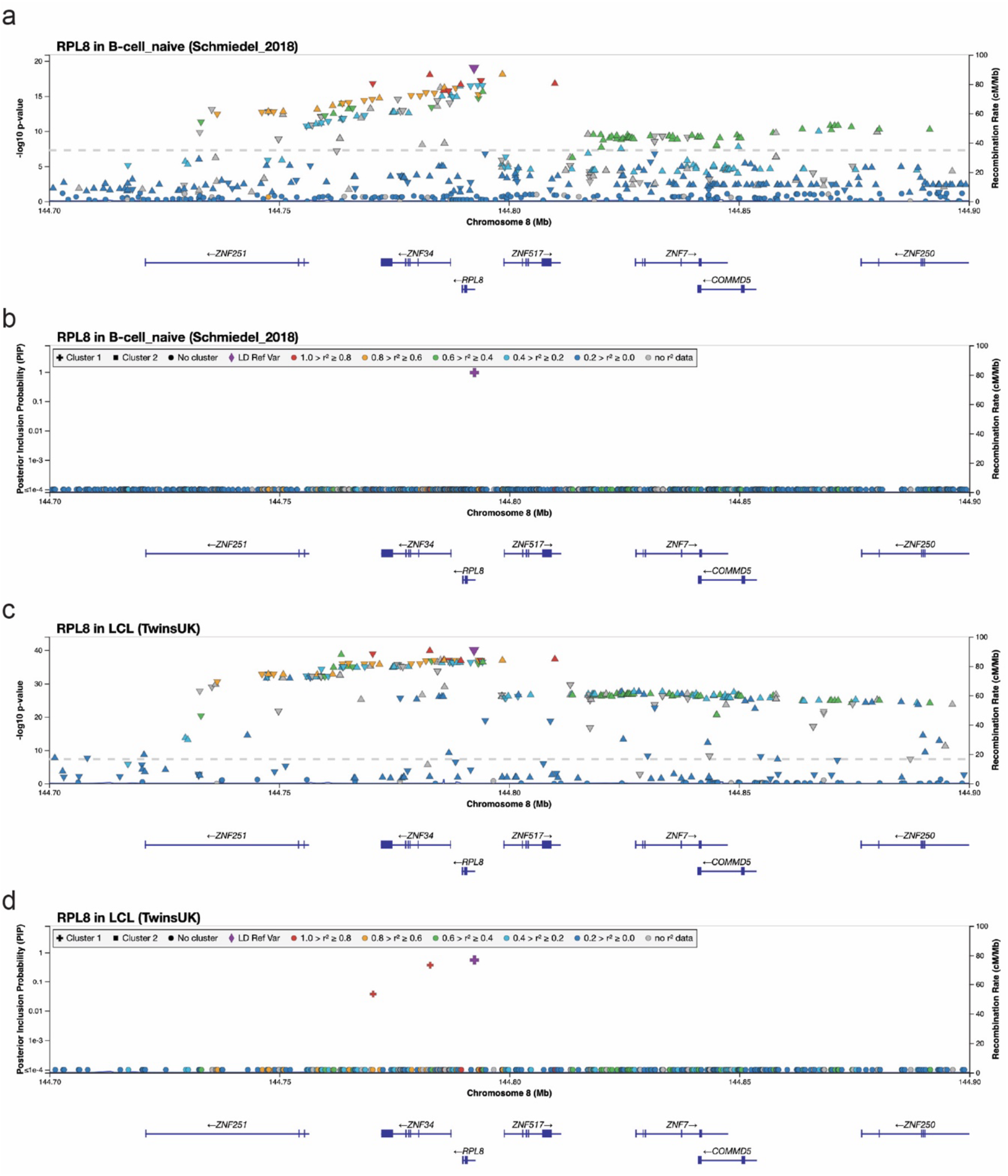
Examination of the RPL8 locus using public data identifies one causal variant. (a-b) Summary of eQTL statistics and posterior inclusion probability from Schmedial et al. Data taken from the fiveX eQTL browser. (c-d) Summary of eQTL statistics and posterior inclusion probability from TwinsUK study. All graphs were downloaded from the publid fiveX eQTL browser. The purple variant is rs2954658 used for CRISPR validation.

**Extended Data Figure 14:**
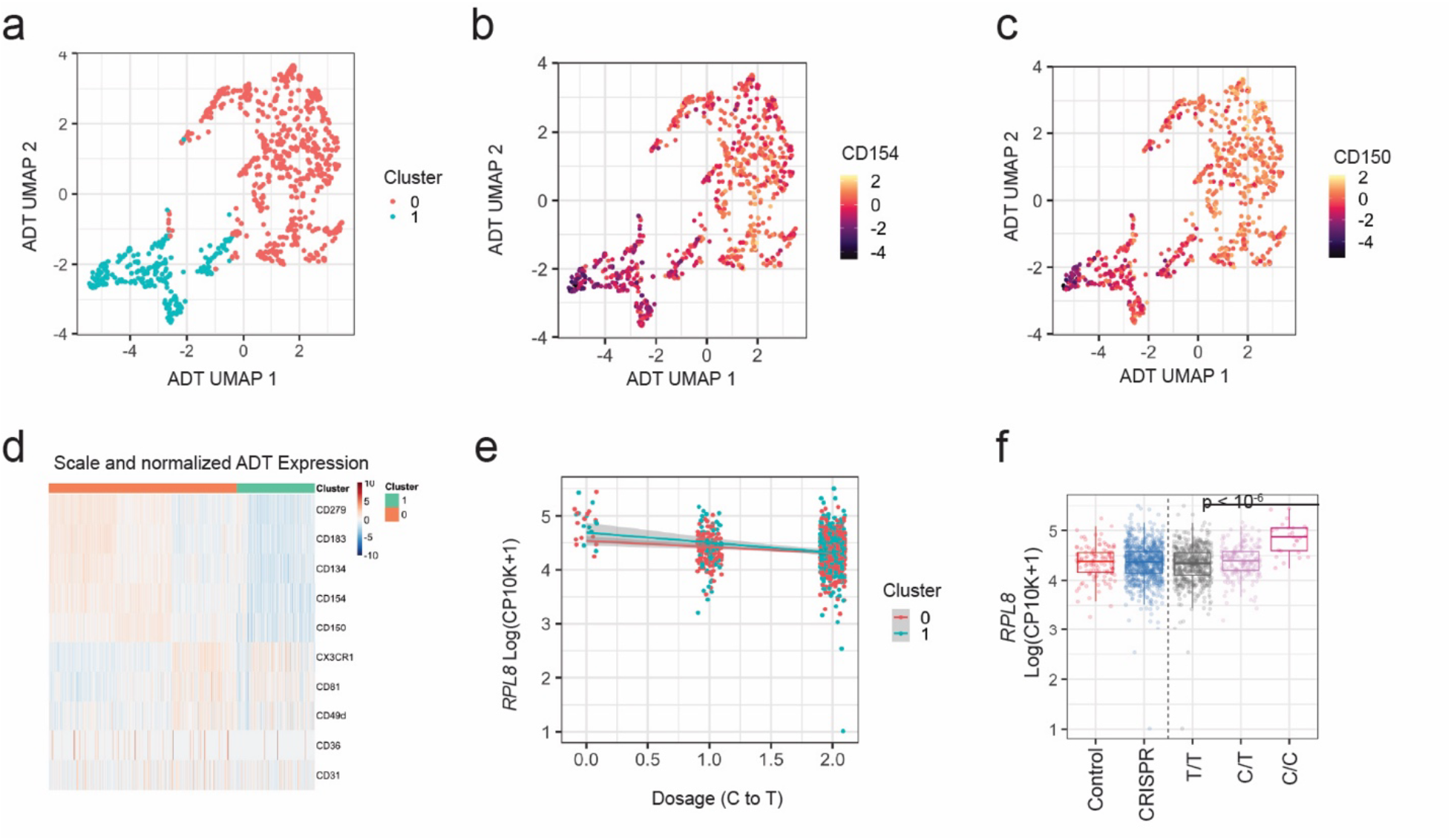
Increase in *RPL8* gene expression is independent of cluster identity. (a) UMAP of all ADTs scaled, normalized, and harmonized by plate identifies two clusters of cells. (b) Scaled and normalized ADT expression of CD154 in ADT UMAP embedding. (c) Scaled and normalized ADT expression of CD150 in ADT UMAP embedding. (d) Heatmap of top variable ADTs, scaled and normalized within each cluster. (e) Normalized gene expression of RPL8 by dosage colored by cluster. (f) Normalized gene expression of RPL8 by dosage and condition. P-values from likelihood ratio tests are plotted.

**Extended Data Figure 15:**
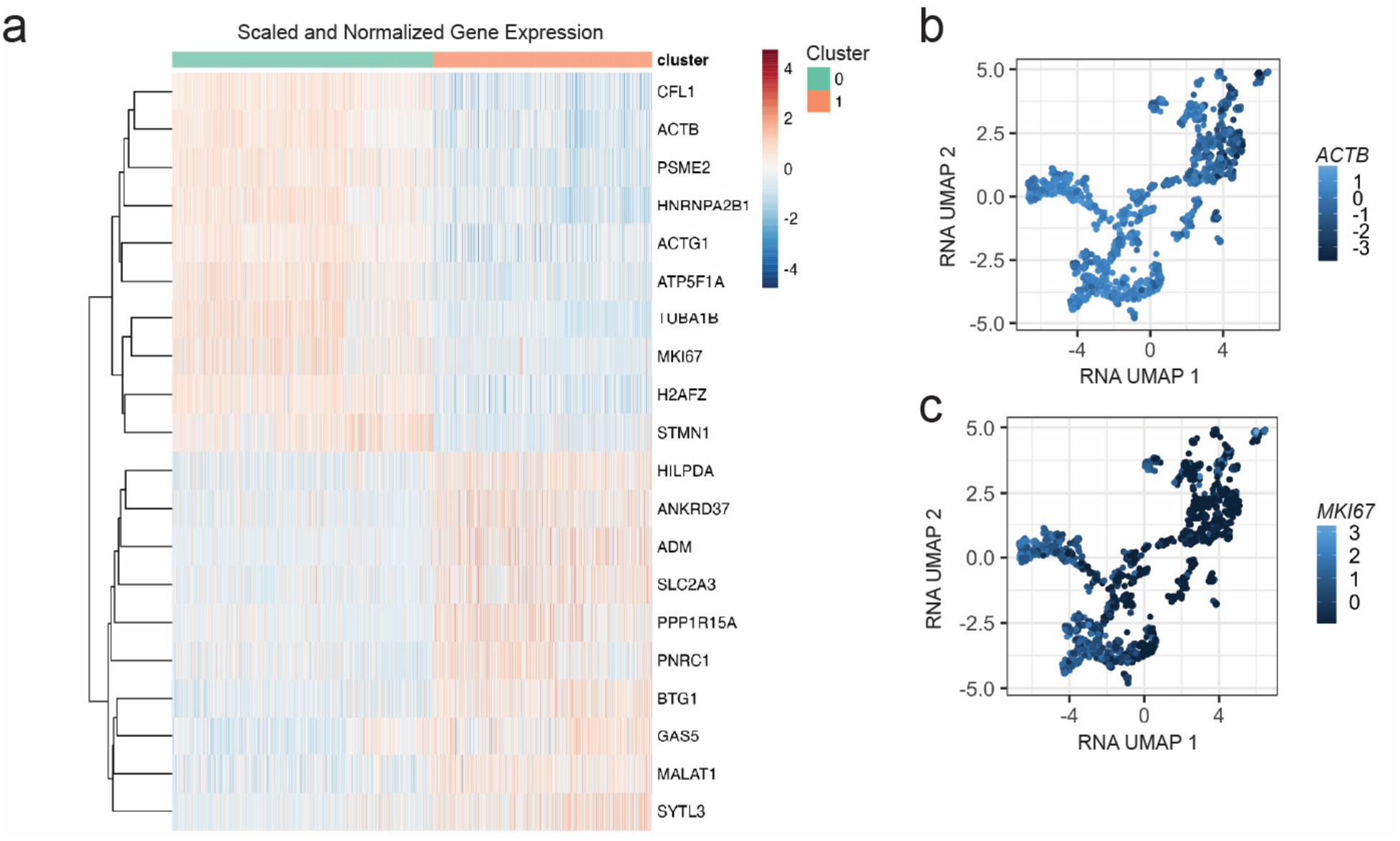
Clustering of *IL2RA* variant edited cells identifies two clusters separated by proliferation status. (a) Heatmap of top variable genes, scaled and normalized within each cluster. (b) Scaled and normalized gene expression of *ACTB* in RNA UMAP embedding. (c) Scaled and normalized gene expression of *MKI67* in RNA UMAP embedding.

## Methods

### Cell line cultures and genomic editing

HH cutaneous T cell lines (ATCC: CRL-2105), Jurkat E6-1 (ATCC: TIB-152), and Daudi cells (ATCC: CCL-213) were cultured in complete RPMI (cRPMI), RPMI 1640 (ThermoFisher) supplemented with 10% heat-inactivated FBS (Rockland), and 1% non-essential amino acids (Gibco), sodium pyruvate (Gibco), HEPES (Gibco), L-Glutamine (Gibco), Penn-Strep (Gibco), and 0.1% b-mercaptoethanol (Gibco). HEK293T cells (ATCC: CRL-3216) were cultured in complete DMEM (cDMEM), DMEM (ThermoFisher) supplemented with 10% heat-inactivated FBS and Penn-Strep antibiotics.

To investigate regulatory regions around *HLADQB1*, HH and Daudi cells were genomically modified as previously described with some modifications^34^. Briefly, 20µM Cas9 protein (Synthego) was mixed with equal volumes of 40µM modified sgRNA (Synthego) and incubated at 37°C for 15 minutes to form ribonuclear protein (RNP) complexes. sgRNA was designed with SpliceR^35^. HH and Daudi cells were nucleofected with 2µL of RNPs in an Amaxa 4D nucleofector and 1ul of ssODN repair template (SE protocol: CL-120 for HH cells, SF protocol: CA-137 for Daudi cells). Cells were immediately transferred to 24 well plates with pre-warmed media and cultured until sorting.

For Jurkat and Daudi cells targeting *FBXO11*, 1µL of mRNA (2µg/µL) encoding the base editor ABE8e-NG was mixed with 1µL of 40µM modified sgRNA (Synthego). Cells were nucleofected with 2µL of mRNA/sgRNA mixture in an Amaxa 4D nucleofector (SE protocol: CL-120 for Jurkat cells, Daudi as above). Cells were incubated as described above in 24 well plates and cultured until sorting.

### Healthy individual recruitment, PBMC isolation, and CD4 T cell magnetic selection

We recruited healthy individuals and processed 40-50mL of peripheral blood under an IRB-approved protocol (IRB# 2008P000427). PBMCs were isolated by layering Ficoll Paque (Sigma-Aldrich) underneath 1:1 PBS-diluted blood followed by centrifugation. Buffy coat layers were extracted, washed in PBS, and then resuspended in cRPMI. Cells were stored by adding an equal volume of RPMI supplemented with 10% DMSO and FBS media and frozen in liquid N_2_ until use. To isolate naïve or total CD4 T cells, frozen PBMCs were quickly thawed and put immediately in warm cRPMI media. Cells were washed twice, and naive or total CD4 T cells were isolated using a magnetic negative selection kit (Miltenyi, Naive CD4^+^ T cell Isolation Kit human, CD4^+^ T cell Isolation Kit) following the manufacturer’s protocols.

### CRISPR Editing of Primary CD4 T cells

After naive CD4 T cell isolation, cells were stimulated with anti-CD3 and anti-CD28 Dynabeads (ThermoFisher) at a ratio of 1:1 (cell:Dynabead) in the presence of 5ng/mL rhIL-2 and 10ng/mL rhIL-7 (Biolegend) in 96 well U-bottom plates. After 1 day, cells are extracted and nucleofected with CRISPR reagents. Briefly, 250-500K stimulated CD4 T cells are nucleofected with 1µL of mRNA (2µg/µL) encoding the base editor and 1µL of sgRNA (40µM, Synthego) in an Amaxa 4D nucleofector (P3 protocol: EH-115). Following nucleofection, cells were transferred to 96 well U-bottom plates and cultured in warm cRPMI for 4 hours. For experiments targeting *PTPRC*, cells were cultured for an additional 5 days with 5ng/mL rhIL-2 before proceeding to staining and cell sorting. For polarization experiments, cells were split into two wells and cultured with either 10ng/mL rhIL-12 (Biolegend), 5ng/mL rhIL-2, and 2ug/mL anti-IL4 (Biolegend) antibody for Th1 conditions or 10ng/mL rhIL-2 and 5ng/mL rhTGFβ (Biolegend) for Treg conditions.

### Cell staining with indexing and oligo-conjugated antibodies

Cells were isolated, washed, and resuspended in a cell staining buffer (2% FBS in PBS with 2mM EDTA). 500K to 1 million cells were stained with fluorophore-conjugated hashing antibodies for 20 minutes on ice. After staining, cells were washed, and conditions pooled together. Pooled samples were then spun down and resuspended in FcX True Stain (Biolegend) (BioLegend) for 5 minutes at room temperature, followed by staining with the TotalSeq-A Human Universal Cocktail 1.0 (BioLegend) for 30 minutes on ice before washing and proceeding to single cell sorting. A full list of antibodies and barcodes can be found in supplementary data.

### Bulk DNA isolation

Cells were pelleted, resuspended in RLT+ buffer (Qiagen), flash frozen on dry ice, and stored at -80°C until processing. For DNA isolation, samples were thawed, vortexed, and incubated for 5 minutes at room temperature before proceeding to DNA isolation using the Qiagen RNA/DNA extraction kit following manufacturer protocols. After isolation, DNA concentrations were measured by spectrometry (Nanovue) and stored at -20°C until use. For analysis of bulk DNA editing, nested PCR reactions were used to amplify 200-400bp genomic regions of interest with custom Illumina adapters added for sequencing. Bulk samples were sequenced alongside single-cell libraries. Sequences of all primers can be found in supplementary data.

### Bulk Flow Cytometry

Stimulated and genomically edited total CD4 T cells were assayed for expression of key protein markers by flow cytometry on day 7 post-nucleofection with a panel of fluorophore-conjugated antibodies. For all samples, cells were isolated, and washed twice in PBS, and FC receptors were blocked with FcX True Stain (Biolegend) for 15 minutes on ice followed by staining with directly conjugated antibodies for 30 minutes on ice. Cells were then washed and samples were analyzed on a BD LSR Fortessa. All data was processed using FlowJo before analysis in R. A full list of antibodies, clones, and manufacturers can be found in supplementary data.

### MINECRAFTseq

Lysis buffer containing 0.2% TritonX (Sigma-Aldrich), 1.2u recombinant RNase inhibitor (Takeda), 1.2mM DTT (Sigma-Aldrich), 1M Betaine (Sigma-Aldrich), 6mM dNTP (Sigma-Aldrich), and 9mM dCTP (ThermoFisher) was added into a 384-well PCR plate using a multi-channel pipette. The Agilent Bravo liquid handling platform was used to add 2.5µM of a well-specific OligoDT barcode into each well, then to distribute 1 µL of barcoded lysis buffer to multiple 384-well plates. These lysis plates were sealed and stored at -20°C until use the next day for sorting. Single cells stained with fluorophore-and oligo-conjugated antibodies were sorted into lysis plates, which were kept sealed on dry ice before being stored at -80°C until use for a maximum of 4 weeks.

For library generation, plates were thawed and incubated at 72°C for 3 minutes, then kept on ice during further processing. 4µL of RT-PCR mix (1.8 uM template switch oligo [Azenta], 0.3mM dNTP, 0.8M Betaine, 4.8mM DTT, 9.7mM MgCl_2_, 0.8U RNase inhibitor, Maxima H Minus Reverse Transcriptase (ThermoFisher), 1x KAPA HIFI buffer, and 2U KAPA HiFi HotStart DNA Polymerase (Roche)) was then added to each well using a MANTIS Liquid Dispenser (Formulatrix). After first strand synthesis (50°C x 1 hour, 85°C x 5 min), a mixture of ADT primers (0.05µM) and genomic DNA specific primers (0.2µM) were added using an I.DOT Liquid Handler (Dispendix) and amplified using the following program: 98°C x 5 min, 18-22 cycles of (98°C x 20s, 65°C x 20s, 72°C x 6 min), 72°C x 5 min, hold at 4°C. After amplification, the samples were split for processing of cDNA, genomic DNA, and ADT.

For genomic DNA: Using the Bravo, 1µL of product per well was taken for further amplification of genomic DNA with nested primers containing a capture oligo with well-specific barcodes. The PCR mix was distributed between plates similar to the process used for lysis plates and contained 0.3mM dNTP, 1x KAPA HiFi Buffer, 2U KAPA HIFI HotStart Taq (Roche), and 0.4µM region specific capture primer (Azenta), 0.4µM region specific P7 primer (Azenta), and 0.4µM capture oligo with well-specific barcode (Azenta). The DNA was amplified using the following program: 98°C x 5 min, followed by 20 cycles of (98°C x 20s, 65°C x 20s, 72 °C x 30s), 72°C x 5 min, hold at 10°C. 2µL of each well containing nested genomic DNA was then pooled using the Bravo and purified with 1.2X solid-phase reverse immobilization (SPRI) beads. Briefly, room temperature SPRI beads were incubated at the given ratio by volume with PCR product for 10 minutes, then placed on a magnet for 5 minutes. Supernatant was removed, and the beads were washed twice with 80% ethanol, dried for up to 15 minutes in air, and resuspended in nuclease-free water (ThermoFisher). The resuspended beads were incubated at room temperature for 5 minutes then placed back on the magnet for 5 minutes, and the supernatant was used for following steps: The cleaned DNA was treated with thermolabile ExoI and ExoVII (NEB) to remove primers (37°C x 30 min, then 87°C x 5 min for ExoI denaturation), and re-cleaned with 1.2X SPRI beads following the same steps as above.

For cDNA, 2µL per well of the original amplification product were pooled using the Bravo and cleaned with 0.6X SPRI beads; the supernatant was saved for ADT sequencing. The cleaned cDNA product was treated with ExoI to remove primers (37°C x 5 min, 85°C x 5 min for ExoI denaturation) and cleaned again with 0.8X SPRI. cDNA concentrations were measured on a QuBit, and 2ng used for tagmentation with the NexteraXT kit (Illumina) per manufacturer’s directions in half volume reactions (25µL total).

For ADT, saved supernatant from the 0.6X cDNA SPRI cleanup was re-cleaned with 1.4X SPRI (2X total) and ExoI treated as above. The ADT product was quantified on a QuBit. For final amplification of pooled samples, 2ng of tagmented cDNA and 5ng of genomic DNA and ADT samples underwent a final amplification step from the 3’ end with custom Illumina-compatible primers. The PCR mix contained 0.2µM P7 and P5 custom oligos (Azenta), 1X Q5 buffer (NEB), 0.5µL Q5 Taq (NEB), 0.2µM dNTP, and nuclease-free water to a total volume of 25µL. For amplification of cDNA, the following program was used: 72°C x 3 min, 95°C x 30s, followed by 16 cycles of (95°C x 10s, 55°C x 30s, 72°C x 30s), 72°C x 5 min, hold at 10°C. For amplification of genomic DNA and ADT, the following program was used: 98°C x 3 min, followed by 16 cycles of (98°C x 15s, 65°C x 20s, 72°C x 45s), 72°C x 10min. After amplification, all samples were re-cleaned using SPRI beads (0.8X for cDNA, 1.2 for gDNA, and 1.6X for ADT) for sequencing. All sequences can be found in supplementary data.

### Illumina Sequencing

Prior to submission to sequencing at the Genomics Platform at the Broad, sample DNA concentrations were quantified on a QuBit, and distributions were measured using a TapeStation (Agilent) with D5000 high sensitivity screentapes. All DNA amplicon libraries were sequenced on a MiSeq for 300 cycles (120 R1, 180 R2). ADT amplicon libraries were pooled at 5% with the RNA libraries and sequenced on a NextSeq550 for 75 cycles (26 R1, 50 R2).

### Processing and alignment of single cell data

I7 and I5 raw FASTQs were merged across different lanes and runs. Transcriptomic reads were then aligned to the human genome (GRCh38 2020-A) with STARsolo (version 2.7.6a)^36^. For all experiments, cells were filtered on at least 500 genes, 1000 UMIs, and less than 10% mitochondrial reads. Counts were log_e_(CP10K+1) normalized and z-score scaled. For dimensionality reduction, PCA was performed on the top 3000 variable genes, as selected using variance stabilizing transformation (VST). This was followed by batch correction with Harmony^37^. Visualization of harmonized PCS was performed with uniform manifold approximation and projection (UMAP) (uwot package)^38^. For index flow cytometry analysis, raw fluorescence values were exported from index sorting as csv files and log_10_ normalized.

Kallisto kite (version 0.27.3) was used to build an ADT reference transcriptome corresponding to the ADT panel (Biolegend TotalSeq A) used and to align ADT reads^39,40^. UMI counts were centered log ratio (CLR) normalized, and z-score scaled. For visualization, PCA was performed on CLR normalized and scaled variable ADTs followed by plate correction using Harmony.

Visualization using UMAP dimension-reduction was performed on harmonized PCs. DNA reads were demultiplexed based on cellular barcodes. Then amplicons were aligned to the amplicon reference using CRISPResso2 (version 2.2.9)^41^ . Allele usage statistics were calculated from CRISPResso2 outputs. For all DNA analyses, cells were filtered on a minimum number of reads (20-40) and minimum allele frequencies (12-25%) on a per-experiment basis. Assuming dizygosity, only the top 2 alleles after filtering were included in the analysis. If after filtering only one allele was recovered, the cell was assumed to be homozygous. Clustering of DNA editing in the DQB1 experiment (Figure 2)was performed with supervised k-means clustering with an n of 3.

### Statistically modeling of gene and protein expression

Associations of gene and ADT expression to dosage and conditions was performed with negative binomial generalized linear modeling. In cell lines we filtered for genes with non-zero expression in greater than 30% of cells and a mean of 2 reads per cell. We modeled counts of gene or ADTs as function of cell-level covariates and fitted a null model 1 using:

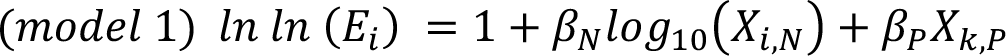

we then compared the above null model to genotype or condition full models using:

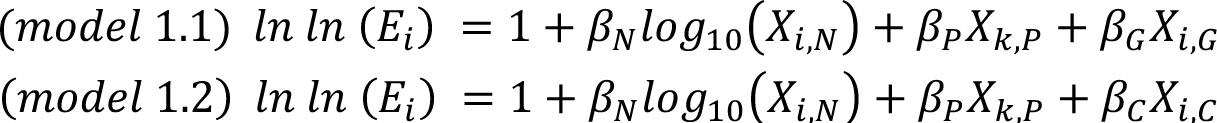

Where 𝐸 is the expected count for cell 𝑖, encompassing gene expression counts or counts associated with ADTs, and all other 𝛽’s represent fixed effects as indicated (N = total number of counts or ADTs, P = plate identity, G = genotype dosage of cell (0,1,2), C = culture condition of cell that is either CRISPR or control) for covariates in cell 𝑖 or plate 𝑘 or donor 𝑑. Specifically, for the DQB1 editing experiment in cell lines we fitted a full model for the deletion size (S) that was calculated in each cell:

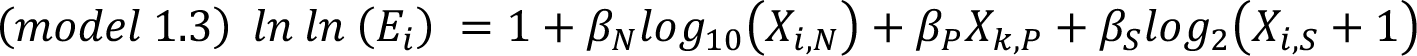

In primary human cells we filtered genes for non-zero expression in greater than 15% of cells. Using a similar negative binomial generalized linear modeling we fitted models with fixed and random effects for primary cells. Specifically, for the PTPRC editing experiments in primary human cells we fitted a null model 2 using:

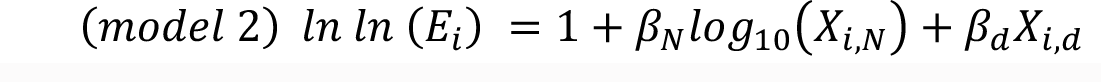

we then compared the above null model 2 using genotype or condition full models:

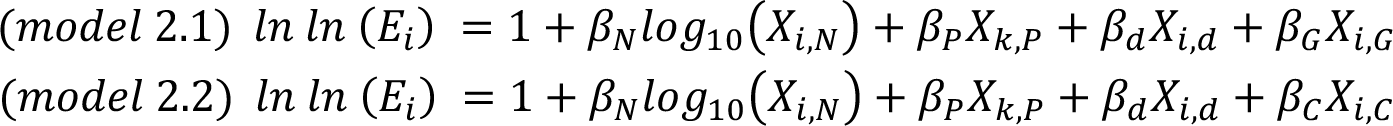

While for the IL2RA editing experiments in primary cells we used random effects for plate and donor. Here we fitted a null model 3 using:

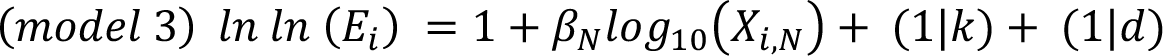

we then compared the above null model 3 using genotype or condition full models:

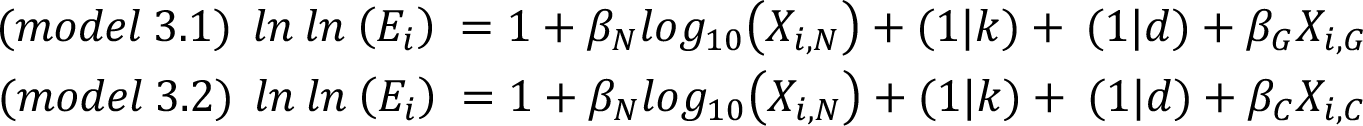

We used the glm.nb function in lme4 (R package) to model fixed effects and glmer.nb function in MASS (R package) model random effects.

To determine the significance of these model, we used a likelihood ratio test (LRT) comparing the null and full models and calculated a p-value for the test statistic against the Chi-squared distribution with one degree of freedom. When indicated, we corrected for multiple hypothesis testing by Benjamini-Hochberg correction (FDR).

### Demultiplexing Donors

Samples were genotyped on the Illumina Multi-Ethnic Genotyping Array (MEGA) or Global Screening Array (GSA). Genotyping quality control, haplotype phasing, and whole-genome imputation was performed for MEGA as previously described^42^. Analogous steps were performed for GSA, with further genotyping quality control including *P*HWE > 1×10^-10^, sample relatedness (IBD) < 0.9, and sample heterozygosity rate > 0.217. All imputed variants were filtered on minor allele frequency (MAF) > 0.01 and imputation accuracy (*Rsq*) > 0.9 in both datasets. Variants were further pruned to the intersection of filtered variants between the two genotyping arrays. Non-biallelic variants and variants within sex chromosomes or non-exonic regions were also removed. Variant coordinates were converted from hg19 to GRCh38 using CrossMap^43^. Demuxlet (version 1.0) was run with default parameters to predict donors from genotypes^44^. Cell donor identity was assigned using the output column “SNG.1ST” corresponding to the most likely singlet identity. Cells with ambiguous donor assignments were excluded.

### Gene set enrichment analysis

We performed gene set enrichment with fgsea^45^. As input, we used the results of the linear model which predicted gene expression from genotype dosage in the CD45 experiment. We included all genes in the enrichment analysis. Genes ranks were defined according to the sign of the effect size of genotype dosage (for directionality) multiplied by the negative log_10_ P-value. We included the MSigDB immunological signatures (C7) pathways^46,47^ that were specific to CD4 T-cells. We then filtered to the pathways that included between 15 and 500 genes.

### Base Editor mRNA

Base editor mRNAs were generated by *in vitro* transcription using the HiScribe T7 High-Yield RNA synthesis kit (NEB Cat No. E2040S) via the method described in^48^. NEBnext polymerase was used to PCR-amplify template plasmids and install a functional T7 promoter and a 120 nucleotide polyadenine tail. Transcription reactions were set up with complete substitution of uracil by N1-methylpseudouridine (Trilink BioTechnologies Cat No. N-1080) and co-transcriptional 5’ capping with the CleanCap AG analog (Trilink BioTechnologies Cat No. N-7113) to generate a 5’ Cap1 structure. mRNAs were purified using ethanol precipitation according to kit instructions, dissolved in nuclease-free water, and normalized to a concentration of 2µg/µL using Nanodrop RNA quantification of diluted test samples.

